# Mutagenesis at non-B DNA motifs in the human genome: a course correction

**DOI:** 10.1101/2022.02.08.479604

**Authors:** RJ McGinty, SR Sunyaev

**Affiliations:** Department of Biomedical Informatics, Harvard Medical School, Boston, Massachusetts 02115, USA; Division of Genetics, Department of Medicine, Brigham and Women’s Hospital and Harvard Medical School, Boston, Massachusetts 02115, USA

## Abstract

Non-B DNA structures formed by repetitive sequence motifs are known instigators of mutagenesis in experimental systems. Analyzing this phenomenon computationally in the human genome requires careful disentangling of intrinsic confounding factors, including overlapping and interrupted motifs, and recurrent sequencing errors. Accounting for these factors eliminates all signals of repeat-induced mutagenesis that extend beyond the motif boundary, and eliminates or dramatically shrinks the magnitude of mutagenesis within some motifs, contradicting previous reports. Mutagenesis not attributable to artifacts revealed several biological mechanisms. Polymerase slippage generates frequent indels within every variety of short tandem repeat motif, implicating slipped-strand structures. Interruption-correcting SNVs within STRs distinctly implicate error-prone Polκ. Secondary-structure formation promotes SNVs within palindromic repeats, as well as duplications within direct repeats. G-quadruplex motifs cause recurrent sequencing errors, while mutagenesis at Z-DNAs is conspicuously absent.

## Main text

By nature of the genetic code, some repetitive DNA sequences are capable of intra-strand base pairing, while other motifs can allow non-Watson-Crick base pairing. This non-canonical base-pairing can stabilize secondary structures, including hairpins, slipped-strand, triplex, Z-DNA and quadruplex structures. Non-B-form DNA structures are disruptive to DNA replication, with possible consequences including polymerase slippage, template-switching, replication fork stalling and collapse, and double-strand breaks. Subsequently, repetitive sequences can cause misalignment during replication restart and homology-directed DNA repair. This can lead to any combination of genomic rearrangements, repeat tract-length changes and single nucleotide mutations within and beyond the motif^1^. Much of this was uncovered in experimental systems, where long repeat tracts model the genetic component of several neurodegenerative disorders.

While the human genome contains numerous repetitive elements, vanishingly few reach long lengths in healthy individuals **(Fig. S1A, Fig. S1B)**. Nonetheless, the number and variety of short repeats in the genome creates an opportunity to investigate mutational mechanisms, by starting with a collection of mutations from large-scale sequencing efforts and measuring their enrichment surrounding repeat motifs. While several groups have performed similar analyses^2,3,4,5,6^, we find strikingly different results after controlling for several key factors: namely, that there is abundant overlap between repeat categories, that motifs are clustered together, that repeats and their flanking regions have highly non-random nucleotide composition, and that repeats are responsible for recurrent sequencing errors. We find that mutagenesis induced by short repeats is limited to a subset of motifs and does not extend beyond the motif boundaries. Polymerase slippage at short tandem repeats (STRs) is the dominant mechanism contributing to this mutagenesis, with structure-forming G4, direct and inverted motifs contributing some additional mutagenesis.

## Results

### Confounders of non-B motif analysis

Sequence symmetry exists along three axes: direct symmetry (a sequence followed by itself), mirror symmetry (a sequence followed by itself in reverse) and inverted symmetry (a sequence followed by its reverse complement). Sequence symmetry is a major component of secondary structure formation: hairpin structures require inverted symmetry, direct repeats support slipped-strand structures, and triplex structures require both mirror symmetry and homopurine sequence content. Certain sequences may contain multiple symmetries. For example, AT di-nucleotide repeats contain direct, mirror and inverted symmetry. Other short tandem repeats (STRs) can also satisfy the requirements of G-quadruplex and Z-DNA motifs. We therefore generated a database of unique repeats by first separating out STRs by motif, and then ensuring that direct, mirror, inverted, Z-DNA, and G4 motifs did not contain overlapping coordinates with any other motif in the database **(Fig. 1, Fig. S1C)**.

**Fig. 1:**
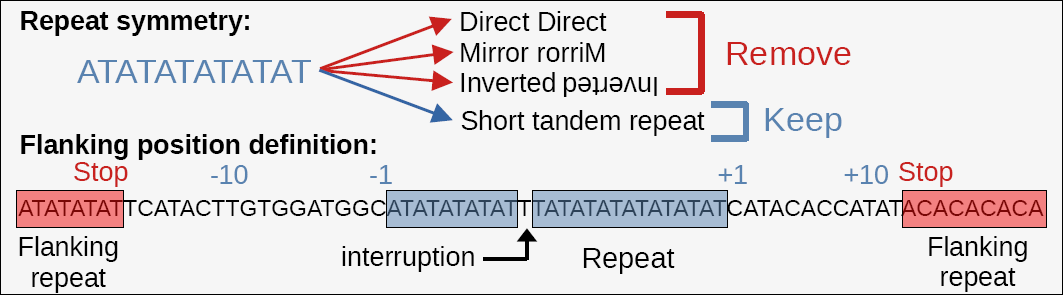
Defining repeat motifs and their flanking positions. Top: example repeat motif containing multiple symmetries, but classified only as an STR. Bottom: Motifs may include interruptions, which are accounted for when defining relative flanking position coordinates. For each motif, the flanking sequence ends when another motif is encountered. Together, this ensures that flanking positions are free of repetitive motifs.

Given the large number of repeat motifs in the genome, many motifs lie in close proximity to one another by chance. In order to avoid mistaking mutations surrounding one motif for mutations that are actually within another motif, we ensured that flanking regions were free of any other discernible repeat motif, including transposable elements **(Fig. 1, Fig. S1D)**. Additionally, we searched for the continuation of a motif past short interruptions **(Fig. 1)**. The advantage is two-fold: interruptions are not mistaken for flanking sequences, and mutations can be categorized by whether or not they perfect the motif.

Repetitive regions are defined by their unusual nucleotide content, but nucleotide composition alone has a large effect on mutation rate. There is a >80-fold difference between the mutation frequency of the lowest and highest trinucleotides due to 5-methylcytosine deamination, and a >10-fold difference between the lowest and highest non-CpG trinucleotide mutations^7^. By normalizing to the observed trinucleotide mutation frequency across the entire genome, we ensure that any observed increase in mutation frequency is not due to the effect of nucleotide composition alone.

We employ the gnomAD database, a large collection of SNVs and indels derived from Illumina short-read genome sequences^8^. Since repeats are known to cause replication errors *in vivo*, we surmised that repeats would likely cause sequencing errors *in vitro*. This is especially likely because the template is single-stranded and free to form secondary structures, and because *in vitro* synthesis lacks a helicase to unwind those structures, as well as other fork-protecting factors. G-quadruplex structures have been shown to induce sequencing errors in both short-read Illumina sequencing and long-read Pac-Bio sequencing, which both employ sequencing-by-synthesis^9,10^. Increasing sequencing depth alone is not a cure-all for this issue, as repeats may induce highly-recurrent sequencing errors^9,11,12^. Furthermore, repetitive regions may also suffer from sequence alignment artifacts. In light of this confounding factor, we employed sequencing quality control metrics to identify mutagenesis signals likely to be affected by sequencing errors. In particular, we see that G-quadruplex motifs and STRs are heavily prone to errors **(Fig 2, Fig S2A, Fig S2C)**.

**Fig. 2:**
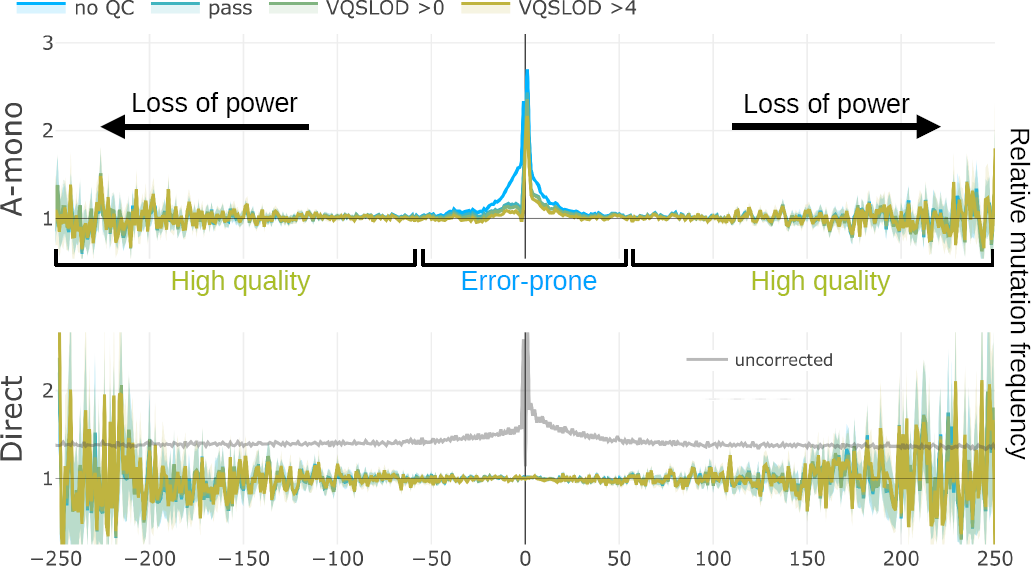
Mutagenesis flanking non-B motifs. X-axes are coordinates relative to central motif (0 position encompasses entire repeat). Y-axes are relative mutation frequency (compared to the gnomAD average, normalized by trinucleotide mutation type). Blue lines: no sequencing quality filters, green and yellow lines: increasingly stringent filters (‘pass’ indicates gnomAD’s passing quality filter based on a VQSLOD score of -2.774). 95% binomial confidence intervals indicated in transparency. Black lines indicate the loss of power at coordinates further away from the central motif due to encountering other motifs. **Top:** A-mononucleotide STR motifs (combined with T-mononucleotide motifs on the opposite strand), demonstrating poor sequencing quality. **Bottom:** Direct repeat motifs, demonstrating comparison to analysis without corrective measures (STRs not excluded, repeats and transposons not removed from flanking positions, no trinucleotide normalization).

### Elimination of repeat-induced mutagenesis at a distance

Experimental studies have shown that long repeats can trigger mutagenesis beyond the motifs, due to the action of break-induced replication (BIR)^1,13^. Thus, we first examined mutation frequencies surrounding repeat motifs. Simply put, we find no evidence for mutagenesis at a distance driven by the short motifs that pepper the reference genome **(Fig 2, Fig. S2)**. Rather, we see that our corrective efforts clearly eliminate false signals of elevated mutagenesis at a distance **(Fig 2, Fig. S2D)**. STRs are internally mutagenic, and their overlap with and proximity to other motifs drives the bulk of spurious signals of mutagenesis at a distance. **(Fig 2, Fig. S2D)**. Signals of excess mutagenesis surrounding G4 motifs and A-mononucleotide repeats are driven by sequencing errors **(Fig. 2, Fig. S2A, Fig. S2C)**.

Because of the large size of the gnomAD database, our method takes into consideration that alleles appearing in more than one individual may sometimes represent independent mutations. We confirmed that this effort eliminates saturation effects by repeating the analysis with a downsampled database **(Fig S2E)**. We further attempted to allay concerns with the quality of the gnomAD database by using *de novo* point mutations gathered from a variety of public sources. This has the advantage of excluding recurrent errors that would be shared between parents and siblings. Unfortunately, despite the relatively large size of our assembled *de novo* database, it was still too underpowered to provide new insights, and some repetitive regions were likely filtered out during *de novo* mutation calling **(Fig S2F)**. At best, we can say that no large increases in flanking mutagenesis emerged from the *de novo* analysis that were missing from the gnomAD dataset **(Fig S2F)**.

For STRs, we see evidence of elevated mutagenesis affecting the immediate upstream and downstream nucleotides, not extending farther than 3 nt **(Fig. 2, Fig. S2A)**. Here we observe an excess of mutations and/or errors that would extend the motif. **(Fig. S2G)**. Because of the difficulty of classifying every potential configuration of an interrupted motif, we know that many motifs in the genome are surrounded by psuedo-motif sequences that escape classification **(see Methods)**. In essence, these regions are not true flanking regions, and what we observe here are interruption-correcting mutations, a phenomenon that we explore further below. The only long-range mutagenic signal that we observed, surrounding CG dinucleotide repeats, is due entirely to their presence within CpG islands, and is not related to the repeat itself **(Fig S2A)**.

Because STRs are known to be potent instigators of indels, we examined the frequency of short insertions and deletions surrounding repetitive motifs. Here the results mostly mirrored those for SNVs, showing the presence of indels in the immediate vicinity of STRs, G4 motifs and direct repeats, but not revealing any long-range mutagenic processes **(Fig S2H)**.

### Mutagenesis within STRs

Having eliminated any elevated signals of long-range mutagenesis, we further examined mutagenesis within repeat motifs. Experimental evidence suggests that a variety of mechanisms contribute to mutations within non-B motifs^1^. In summary, we find that only a subset of motifs are mutagenic within the motif, while sequencing errors are frequent.

We measured the rate of SNVs and indels within STRs with respect to the motif length. We separately measured mutations within perfect motifs and those with interruptions, differentiating between mutations that remove imperfections and mutations that introduce imperfections. We found that indels were highly elevated within STRs in a motif length-dependent manner **(Fig. 3A, Fig. S3A)**, as would be expected under the polymerase slippage model. Consistent with previous findings^14^, we see that most STRs are biased towards contractions at lower motif lengths, switching to an expansion bias at longer motif lengths **(Fig S3C)**.

**Fig. 3:**
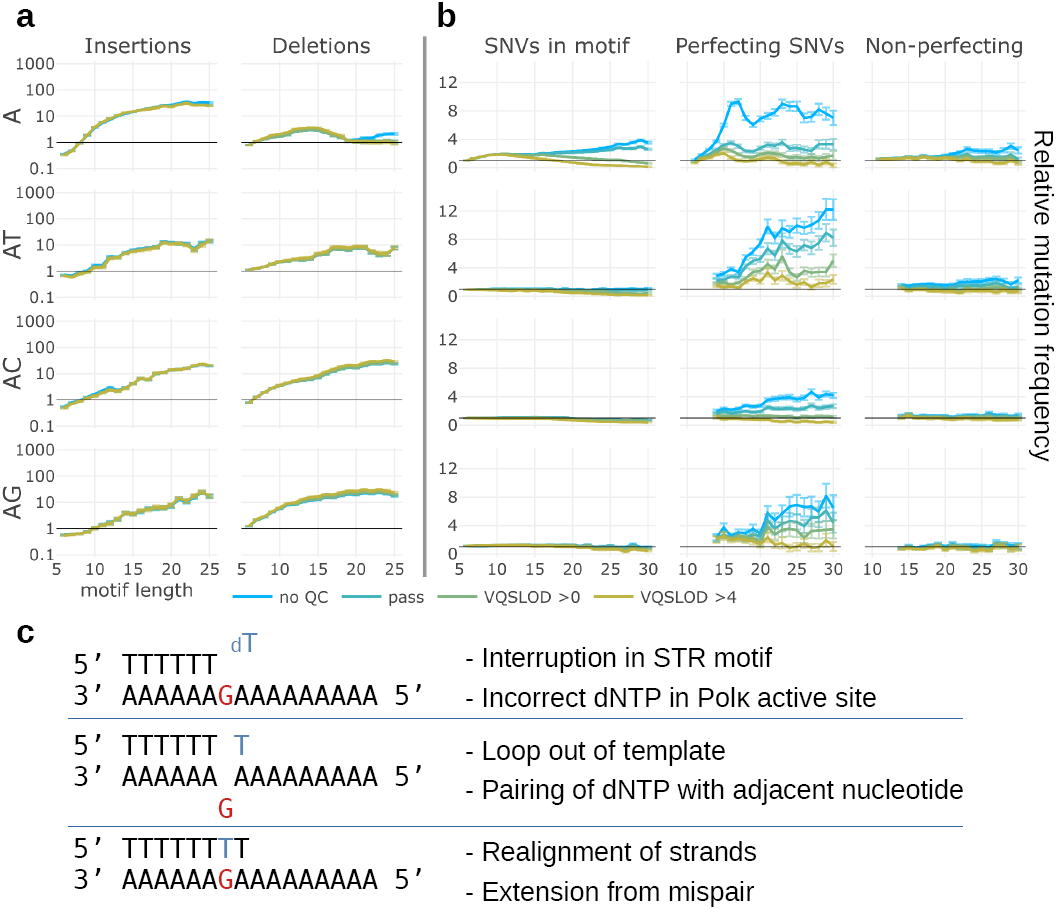
Polymerase slippage generates mutations, indels and sequencing errors at STRs. Y-axes are relative mutation frequency. Error bars indicate 95% binomial confidence intervals. X-axes are motif lengths. Blue: no sequencing quality filters, green and yellow: increasingly stringent quality filters. Motif sequence (combined with their reverse-complements) indicated at left of each row. **a)** Insertions and deletions within STR motifs. **b)** SNVs within perfect motifs, and SNVs that perfect or don’t perfect interruptions within imperfect motifs. **c)** Model for interruption-correction by Polκ during rescue from polymerase pausing.

We also observe elevated rates of SNVs within STRs, though the relative increase is smaller than for indels. For all STRs, we observe SNVs that perfect interruptions within STRs **(Fig 3B, Fig. S3B)**. (As discussed above, substitution types at immediate flanking positions strongly suggest that these mutations should also be considered interruption-perfecting.) The rate of this phenomenon generally increases with longer motif lengths, as does the rate of sequencing artifacts. While errors are frequent, enough SNVs remain after stringent quality filtering that this phenomenon must also have a biological basis. Note that such errors would cause under-counting of legitimate novel mutations in perfect STRs. This effect should be proportional to the observed rate of errors at known STR interruptions in the reference genome, resulting in modest numbers of missed SNVs within STRs.

Strikingly, we see a sharp contrast between mononucleotide and higher-order STRs in the rate of SNVs that would disrupt the motif **(Fig. 3B**, **Fig. S3B)**. In mononucleotide repeats, we observe an approximate balance between mutations that perfect and disrupt the motif. In contrast, di- and trinucleotide STRs show elevated rates of perfecting mutations, but the rate of disrupting mutations does not rise above the background mutation rate. This raises the possibility that disrupting mutations in mononucleotide repeats arise from a mechanism other than polymerase slippage.

Complicating this picture, we also measured the fidelity of STR expansions. We categorized insertions within STRs into those that created a perfect expanded repeat, those that generated an expansion with 1-2 interruptions, and those that inserted an unrelated sequence. Imperfect expansions are assumed to result from low fidelity during synthesis of the insertion. By summing the total number of single base errors within expansions and comparing to the total number of bases inserted, we derived the per-base density of synthesis errors during expansion, which ranges from ∼10^−3^ to ∼10^−2^ errors per nt, differing by STR sequence and motif length **(Fig S3D)**. This is in stark contrast to the low rate of motif-disrupting SNVs when a length change is not observed.

The question also arises as to how polymerase slippage generates interruption-perfecting SNVs. We have already noted the very high relative rate of indels within STRs. Deletion of an interruption, along with a length-compensating insertion elsewhere in the motif, would be indistinguishable from a motif-perfecting mutation, similar to an effect previously reported at compound STRs^15^. To estimate the likelihood of these two events occurring independently, we compared the absolute frequency of SNVs and indels within STRs **(Fig. S3E)**. While the absolute frequency of indels is often two orders of magnitude greater than SNVs, the product of the frequencies of an insertion within the motif and a deletion of the interruption would be 2-3 orders of magnitude lower than the frequency of an interruption perfection. Thus, this result calls for an alternative model that occurs in a single event.

### Mutagenesis within other non-B motifs

We further investigated SNVs and indels within direct, inverted, mirror, ZDNA and G4 motifs.

Direct repeats showed limited signs of elevated SNV mutagenesis, but exhibited a peculiar pattern of indels. We observe only a small increase in SNVs immediately flanking the motif, which scales with increasing motif length and decreasing spacer length **(Fig. S4A)**. We observed a much greater relative increase in the rate of indels. In particular, we observed deletions within the motif, as well as insertions that appeared at the beginning of the motif and at interruptions **(Fig. S4B)**. Repeats with spacers under 10 nt exhibited the most insertions **(Fig. S4B)**. We further saw that only insertions longer than 5 nt were elevated **(Fig 4A, Fig. S4B)**. We confirmed that these long insertions represent partial duplications of the motif, typically encompassing one of the two repeats along with the spacer sequence **(Fig. S4C)**. Interruptions appear to influence the start and stop of the duplication, appearing at the insertion boundaries more frequently than expected **(Fig. S4D)**.

**Fig. 4:**
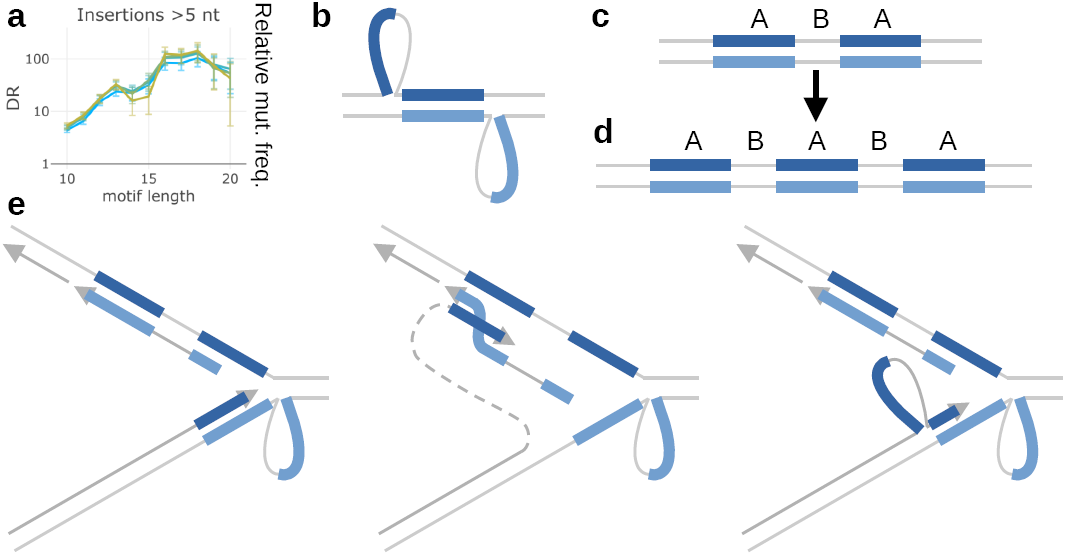
Duplications at direct repeat motifs. **a)** Insertions of >5 nt within direct repeats. Y-axis is relative mutation frequency. Error bars indicate 95% binomial confidence intervals. X-axis is motif length (length of one repeat, not including spacer). Blue: no sequencing quality filters, green and yellow: increasingly stringent quality filters. **b)** Direct repeats adopting a slipped-strand secondary structure. Loops include one copy of the repeat and the spacer sequence. Loops may be stabilized by *ad hoc* basepairing (not pictured). **c)** Direct repeats (A) separated by a spacer sequence (B). **d)** Frequently observed A-B-A-B-A arrangement following duplication of motif and spacer. **e)** Model for motif duplication during replication. Left: leading strand pauses after encountering secondary structure. Middle: Template switch of the leading strand to copy from the nascent lagging strand Okazaki fragment, which may begin within the second copy of the repeat. Right: After replicating to the end of the Okazaki fragment, the nascent leading strand switches back to the leading strand template, landing again within the first repeat. Upon completion of replication, the resulting arrangement would be A-B-A-B-A.

We further investigated the fidelity of duplications within direct repeats. Of 11,869 insertions with a length of 10 nt or greater, we found 5,382 where the insertion perfectly matched a region within the motif, 550 with a single base mismatch, and 79 with two mismatches. Treating these mismatches as polymerase errors during synthesis of the duplication, we calculated the error density to be ∼2×10^−3^ errors per nt, with a spectrum favoring transversions, when compared to the spectrum of *de novo* germline mutations **(Fig. S4E)**. We also found 33 dinucleotide mismatches, representing an error density of ∼3×10^−4^, and with a spectrum that favored NN>TT or NN>AA mutations **(Fig. S4F)**.

In contrast to direct repeats, inverted repeats appear prone to SNVs rather than indels **(Fig. S5A, Fig. S5B)**. While issues with low power obscure the analysis, one clear pattern emerges: inverted repeats separated by spacers of 1 or 3 nt appear to be excessively mutagenic at the center of the spacer **(Fig. S5A)**, similar to a previous report^3^. This occurs in a stem length-dependent manner. Though power-limited, we don’t see a strong elevation of APOBEC-related TCN>T mutations at these spacers **(Fig S5C)**. We see that spacers of 2 or 3 nt more frequently mutate to extend the inverted motif, removing or shortening the spacer, respectively **(Fig. S5A)**. We find no evidence that the positions flanking IRs frequently mutate in a manner that would extend the motif **(Fig. S5A)**. Thus, inverted symmetry would tend to extend inward to shrink the spacer, but not outward into the genome, limiting self-propagation of symmetry. While power-limited, we see indications that IRs with short spacers may also be mutagenic within the stems of the motif, more frequently mutating to correct interruptions **(Fig. S5A)**.

We did not observe any convincing signals of excess mutagenesis within mirror repeats, whether SNVs or indels **(Fig. S6A, Fig. S6B)**. However, it should be noted that homopurine mirror repeats, which may form H-DNA structures, are very rare in the genome following exclusion of STRs, precluding their separate analysis. Mirror repeats with heterogeneous nucleotide content are not known to form any non-B structures, so the lack of elevated mutagenesis is the expected result.

A more surprising negative result is that for Z-DNA motifs. We observe no elevation of SNVs or indels within Z-DNA motifs **(Fig. S6C)**. Prior to our filtration of STRs, AC/TG dinucleotide repeats made up a large portion of the Z-DNA motif category. Mutagenesis at AC repeats is similar to other dinucleotides. Together, this suggests that Z-DNA formation not a contributor to mutagenesis in the human germline.

Finally, we observe that G-quadruplex motifs are instigators of recurrent sequencing errors, especially T>G mutations on the 5’ end of spacers (sequences between G-runs) **(Fig 5)**. After accounting for errors, G-runs appear mildly mutagenic, while spacers do not **(Fig 5)**. This refutes earlier reports that single-strandedness predisposes spacers to mutation. The situation for indels is slightly different. Spacers again appear to trigger erroneous insertions and deletions **(Fig. S7A, Fig. S7B)**. However, we also see elevated levels of high quality indels within G4s, with spacers biased towards insertions **(Fig. S7A)** and G-runs biased towards deletions **(Fig. S7B)**.

**Fig. 5:**
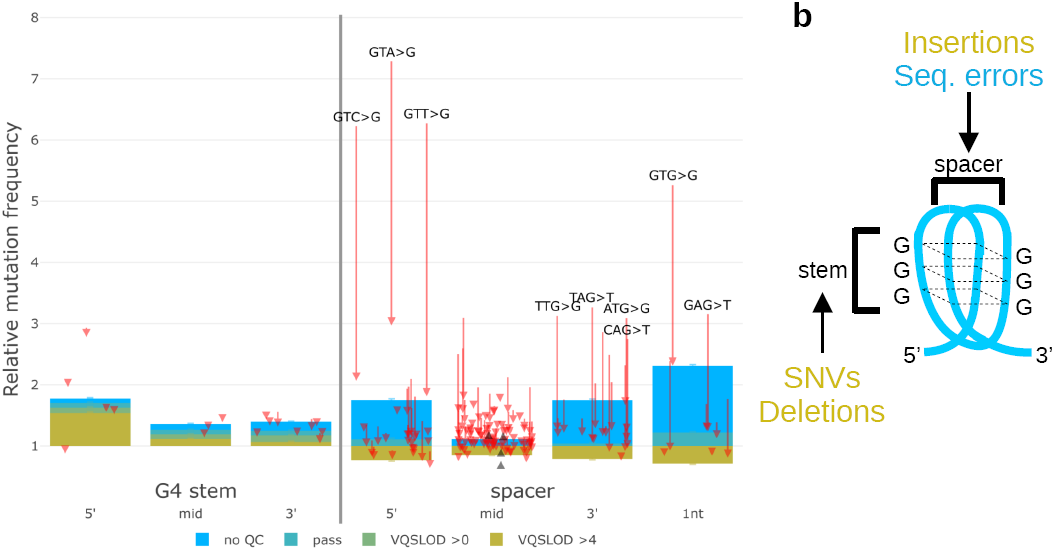
G4 motifs are prone to recurrent sequencing errors. **a)** Mutations within G4 motifs. X-axis indicates positions within G-runs, or spacers between G-runs. Spacers of 1 nt in length are separated. Y-axis is relative mutation frequency. Blue: no sequencing quality filters, green and yellow: increasingly stringent filters. Error bars indicate 95% binomial confidence intervals. Arrows indicate magnitude and direction of change for individual trinucleotide mutation frequencies before and after sequencing quality filters. Mutations with large magnitude changes are highlighted in text. **b)** Diagram of a G-quadruplex structure. Spacers are prone to insertions and sequencing errors, while G-runs are prone to SNVs and deletions (see Fig. S5).

## Discussion

Ultimately, after carefully avoiding issues confounding the analysis, we find that short repeat motifs in the human reference genome pose little mutagenic threat to the surrounding genome. All signals of elevated mutagenesis surrounding short repeat motifs are eliminated after carefully removing confounding factors. This suggests that DNA replication and repair systems are generally capable of handling short repetitive sequences, with little more than polymerase slippage as a consequence. These results are actually in line with experimental systems, which demonstrate a repeat length threshold for large-scale instability.^1^ Short repeats are mostly stable, undergoing small-scale expansions and contractions. Completely derailing replication requires very long repeats, leading to extreme outcomes such as BIR. While experimental work demonstrates that long-range mutagenesis is a consequence of BIR, the more detrimental consequence is that of frequent genomic rearrangements^16^. For this reason, repeats capable of triggering BIR cannot be sustained for long in the germline, and thus are very rare in the genome and in the population **(Fig. S1A, Fig. S1B)**.

In contrast, short motifs are relatively stable. The frequency of SNV mutagenesis within motifs does not stand out in comparison with other mutagenic processes that are active in the human germline, including those driven by sequence context^7^ and regional variation^17^. Thus, short motifs are not major sources of SNVs contributing to population diversity or common disease risk. SNVs in STRs tend to correct mismatches, ensuring the continued presence of the motif. Because short motifs are not destructive to their surroundings, this resolves some tension in the notion that non-B DNA structures may serve functional roles in biology^18,19,20,21^. Indeed, patterns of indels at G4 sequences would tend to lengthen the spacer and shorten the G-runs, leading to the loss of structure-forming potential, suggesting that G4 motifs must be maintained evolutionarily.

As we and many others have observed, even short STRs are highly prone to length alterations via polymerase slippage. Expansion and contraction of STRs may indeed contribute to population diversity, expression of numerous genes, and complex traits identified by GWAS^22,23^. The question remains as to the life cycle of STRs in the genome. We also confirm prior observations that many STRs are biased towards contractions at shorter motif lengths and expansions at longer motif lengths^14^. STRs are exponentially more common in the genome at shorter lengths, suggesting that the vast majority of STRs will tend to remain very short. Our results both clarify and complicate previous work suggesting that as STRs grow, they will increasingly accumulate disruptions to the motif, preventing further expansion^6^. Only for mononucleotide repeats do we observe an increase in motif-disrupting SNVs. For all other STRs, interruptions appear to be gained only during the expansion (and possibly contraction) process due to low-fidelity synthesis (in addition to interruptions due to the background mutation rate). However, we see that longer STRs increasingly correct these interruptions, effectively lengthening the motif. Thus, it appears as though STRs that begin expanding can continue to do so until limited by the constraints of natural selection.

We found evidence for several mutational mechanisms operating at non-B motifs, most notably polymerase slippage. Frequent indels at STRs are the hallmark of polymerase slippage, as secondary structures disrupt synthesis while the repetitive sequence makes realignment likely to occur out of register^1^. We see elevated rates of indels at all STR motifs which we had the power to analyze, suggesting the importance of slipped-strand structures, since that is the only structure common across all STR sequences **(Fig. 3A, Fig. S3A)**. This was a surprising finding, as much of the prior literature links STR instability to the peculiarities of each motif and its accompanying non-B structure^24^.

Previous experimental findings indicate that replicative polymerases make few substitution errors at STRs, compared with specialized polymerases involved in DNA repair^25^. This dichotomy may explain why we observe a high density of single-base errors made in the process of repeat expansion, but we see SNVs only at the background mutation rate. When polymerase slippage occurs, it is likely that a low-fidelity polymerase reinitiates synthesis on a misaligned template; in the absence of slippage, high-fidelity synthesis proceeds in-register.

Mononucleotide repeats appear to represent a special case. While they clearly trigger polymerase slippage, we also observe a unique increase in interruption-generating SNVs not elevated within di- and trinucleotide repeats. Bacolla and colleagues proposed that such mutations could be linked to the flexibility of the DNA strand, as dictated by interactions between neighboring bases^26^. Indeed, we replicate the finding that the first and last nucleotide of A/T-tracts are the most mutable, while C/G tracts are more mutable further within the tract **(Fig S3B)**, supporting this model. However, from our data it appears that this mechanism quickly loses relevance for motifs with a longer unit length.

We found that the rate of interruption-correcting SNVs in STRs could not be explained by the length-neutral combination of two independent slippages, and thus that this type of SNV arises from a single event. (It is possible that a high rate of indels during *in vitro* synthesis, combined with pre-phasing errors^27^, alignment errors and other sequencing artifacts, could explain the frequent low quality SNVs of this type.) As discussed above, polymerase slippage can result in the recruitment of low-fidelity polymerases. The unique properties of Polκ, as demonstrated *in vitro*^28^, make it the perfect candidate for such a mechanism **(Fig 3C)**. When provided an incorrect dNTP, Polκ can accommodate a small loop out of the template DNA, allowing it to correctly pair this dNTP with a neighboring nucleotide. Polκ will next realign the template and nascent strands, and extend synthesis from the mispair. This has the effect of swapping a potential deletion for a misincorporation, which is beneficial in the context of avoiding frame-shift mutations. This is strikingly similar to the outcome that we observe here, where interruptions in STRs consistently revert without a change in length.

We observe several mechanisms at play in non-STR motifs. Double-strand break repair by single-strand annealing (SSA) is well known to cause deletions between direct repeats^1^. More surprising is the elevated rate of duplications between direct repeats, which are not a consequence of SSA. Duplications usually include one copy of the repeat and extend to the end of the spacer, such that the initial A-B-A **(Fig 4B)** arrangement would become A-B-A-B-A **(Fig 4D)**. The rate of duplications is influenced by stem length, loop length and the presence of interruptions, which suggests that the stability of slipped-strand structures **(Fig 4A, Fig. S4B)** is an important component. We also found the fidelity during synthesis of duplications to be in the range of low-fidelity polymerases^29^. We can imagine two different mechanisms that would generate duplications in this manner. In the first **(Fig 4E)**, replication encounters a slipped-strand structure, which is stabilized by *ad hoc* basepairing within the loop (which potentially pulls in interrupted portions of the repeat). Leading strand synthesis pauses after replicating the first repeat, while on the lagging strand an Okazaki fragment begins partway within the second repeat and is able to proceed through the first repeat. In this case, the leading strand could use template-switch to copy from the nascent lagging strand until reaching the end of the Okazaki fragment and switching back to the leading strand template. Landing again within the first repeat would generate the A-B-A-B-A pattern. This is similar to a mechanism responsible for large-scale repeat expansion in STRs^1^, and distinct from another mechanism proposed to explain A-B-A-B-A duplications in *E. coli*^30^. Another interesting possibility is that the unpaired regions of the slipped-strand structure may be subject to post-replicative gap-filling **(Fig. S4G)**, a pathway known to involve error-prone synthesis by Rev1/Polζ^31^. We also observe a spectrum of infidelity favoring transversions, consistent with synthesis by Polζ^32^. Furthermore, we observed a high density of double nucleotide substitutions with a spectrum favoring NN>AA or TT, similar to the known GC>AA or TT signature of Polζ^32,33^. Gap-filling of slipped-strand structures would result in pseudo-Holliday junctions, which could be resolved to produce the A-B-A-B-A duplication pattern **(Fig. S4G)**. Thus, post-replicative gap-filling by an error-prone polymerase is a plausible mechanism for the observed duplications.

Elevated SNVs within inverted repeats may be explained by several mechanisms, though polymerase slippage is ruled out by the lack of accompanying indels. Mutagenesis is dependent on stem and spacer length, and thus on the stability of the hairpin structure. Most mutagenesis is concentrated within the unpaired spacer sequence between repeats. While others have suggested a role for APOBEC^3^, which deaminates single-stranded DNA, we don’t see strong evidence of the related TCN>T mutation spectrum **(Fig S5C)**. Mutagenesis is most elevated in extremely short 1-3nt spacers, rather than long spacers which expose more single-stranded DNA, which may instead suggest that helical stress at the bend of the hairpin is a factor. Because we see an elevated rate of mutations that extend the motif into spacers (and possibly at interruptions in the hairpin stem, though the data is power-limited), this implicates either of two pathways: mismatch repair and/or template-switch during replication. An interruption within a stable hairpin stem would appear identical to any other mismatch; a 2-3nt hairpin capped-end is a possible but less obvious substrate for mismatch repair **(Fig S5D)**. The alternate possibility, template-switch **(Fig S5E)**, is well-studied in bacterial model systems, and there is evidence that it also operates in humans^34,35,36^. Interestingly, template-switch mutations are difficult to predict using models that account only for the sequence context of adjacent nucleotides, because the mechanism requires symmetry rather than nucleotide content for both the context and the outcome of the mutation. In any event, the elevation in mutagenesis above the background rate is modest. Indeed, in disentangling STRs from other repetitive motifs, we have generally shown that a motif’s status as a short tandem repeat far outweighs other considerations.

The lack of contribution to mutagenesis from Z-DNA motifs was surprising, given the number of reports linking Z-DNA motifs to genomic instability^37,38^. One possibility is that, where many past studies have used CG or TG dinucleotide repeats as the stereotypical Z-DNA-forming motif, we classify these first and foremost as STRs. We examined two frequently used Z-DNA motif definitions, one specifying RY (purine-pyrimidine) repetitions excluding AT dinucleotides, and the other further restricting to GY repetitions. Because we filter for uniqueness among all of the potential non-B categories, it could be that Z-DNA formation requires some additional property such as mirror symmetry. Another explanation is that our study focuses on mutagenesis in the human germline, whereas most other studies have examined cancer genomes, cultured cells or model organisms. Excitingly, a recent study characterized a zinc-finger protein, ZBTB43, which removes Z-DNA structures^39^. This enzyme is highly expressed for a brief window in mouse germline development, where it facilitates epigenetic reprogramming at CG-rich sequences. Thus, we may observe a lack of Z-DNA-related mutagenesis in the germline because Z-DNA structures are efficiently removed. It is certainly possible that additional structure-specific enzymes, including various specialized helicases^1^, contribute to the limitation of non-B DNA-related mutagenesis in the germline.

## Methods

### Variants

Human germline variants were downloaded from the gnomAD consortium^40^, with version 3.1 consisting of 759,302,267 variants derived from 76,156 whole genome sequences. We filtered this set to single nucleotide variants (SNVs) with a maximum allele frequency of 0.01%. Germline variants of sufficient rarity have previously been shown to serve as a good proxy for mutations, since their composition has not been detectably altered by selective pressure^7^. gnomAD provides a recommended quality cutoff based on allelic imbalance and allele-specific VQSLOD, which is a machine-learning-derived sequencing quality metric based on a combination of strand bias, read position, mapping quality, and read depth^8^. gnomAD’s passing VQSLOD score is -2.774 and above, which excludes 12.2% of SNVs, and we applied additional filters above VQSLOD 0 and 4, where indicated. Rather than deciding on a singular quality cutoff, which would necessitate a trade-off between over-filtering some sites and under-filtering others, we present mutation frequencies without quality control, and across this range of quality filter regimes. This highlights regions prone to potential sequencing quality artifacts (where the filtered and unfiltered frequencies are distinct), and demonstrates that mutagenesis in other regions is invariant to sequencing quality (where the filtered and unfiltered frequencies overlap).

Because of the large number of genomes in the gnomAD database, we considered that some loci with an allele count >1 could be produced by independent mutations in multiple individuals. However, shared ancestry is the primary contributor to higher allele counts. We employed a model^17^ to estimate the number of independent mutations for a given allele count. This model estimates the number of recurrent mutations at a site as a logarithmic function dependent on allele count, the number of sequenced genomes in the dataset as well as the mutation rate at each given site. Since we do not know the mutation rate *a priori*, we assumed a mutation rate 50-times the baseline genome mutation rate. This results in a logarithmic curve predicting on average ∼1.4 mutations for an AC of 2, with a maximum of ∼1.8 mutations for AC values above 6. The mutation rate assumption was conservative to avoid underestimating recurrence, as earlier iterations of the current study suggested that the mutation rate at our sites of interest would not exceed 50-times the baseline mutation rate.

We also believe that overestimation of recurrence is minimal, both because the gnomAD database is not fully saturated, and because lower values of the mutation rate term result in only a slightly lower maximum correction factor. To further study the effects of saturation, we applied downsampling to the gnomAD database. Because the gnomAD database is de-identified, we could not check for saturation effects by downsampling to remove individuals. Instead, we proportionally downsampled the database per each allele count bin, using the downsampling fraction multiplied by the allele count. We further applied the allele count correction to the downsampled database, after adjusting for reduced sample size.

The gnomAD database also includes 44,056,957 indels. VQSLOD values have a different distribution than for SNVs. gnomAD chooses -1.0607 as the passing cutoff, and we additionally chose filter values of 0 and 1.4, in order to mirror the proportion of variants filtered in the four cutoffs for SNVs.

We also gathered *de novo* point mutations from a variety of publicly available sources^41,42,43,44,45,46,47,48^. These studies confirmed the *de novo* status via comparison to parental sequences (ie. trios, quads, and some extended pedigrees). In total, we gathered 617,588 point mutations from 10,912 whole genome sequences, or ∼56 mutations per genome. We converted all coordinates to hg38, where applicable.

### Repeat motif database generation

Where indicated, repeats motif coordinates were gathered from Non-B DB^49^. However, because this database does not account for interrupted motifs, we generated our own database of repeat motifs. Starting from the hg38 human reference genome assembly, we first masked transposable elements based on Repeatmasker ‘non-simple repeat’ regions, both downloaded from the UCSC Genome Browser^50^. We searched autosomal chromosomes for all non-redundant STR motifs from 1-9 nt, using custom Python scripts employing ‘regex’ matching. Coordinates (consisting of chromosome, start position and end position) were then extended upstream and downstream to search for continuation of the motif, including partial motif extensions. For each motif, coordinates were then reduced to a minimal set of non-overlapping coordinates, allowing for up to one interrupted position on either side. The complete set of STRs was then used to mask the reference genome prior to all additional ‘regex’-based searches.

Importantly, the initial motif search required a choice in minimum length requirements. We used a minimum of 5 units for mononucleotides, 3 units for dinucleotides, and two units for trinucleotides and above. Longer length requirements were used, where noted, following the incorporation of interruptions. We required that interruptions must link two segments that each meet the minimum length requirements. Thus, longer length requirements make the motif more easily recognizable., while shorter lengths maximize detection of interrupted motifs. However, choosing length requirements too short would undermine the classification of motifs. For an A-mononucleotide repeat, for example, it is not clear *a priori* whether 1-4 As following an interruption is relevant to mutagenesis. Similarly, it is not clear whether interruptions of 2 or more nucleotides should be allowed for any given motif. Thus, while our method captures many of the possible motif configurations, capturing all is an impossibility, and some flanking regions may still be misassigned. Based on a 5 nt minimum length requirement, most of this uncertainty should exist within 5 nt surrounding the motif.

Pairs of motifs containing 5 nt of mirror or inverted symmetry separated by no more than 100 nt were found via ‘regex’ searches, and then coordinates for each pair were extended both outwards and inwards until symmetry was lost, or a masked region was reached on either side. 1 nt interruptions were then allowed on either side, and coordinates were again extended both outwards and inwards until symmetry was lost. For direct repeat motifs, the search proceeded left from the 5’ ends of both motifs, and right from the 3’ ends, again allowing for a single interruption on either side. Because the left/right extension necessarily shrinks the spacer distance, the initial search parameters allowed pairs to be separated by up to 1000 nt, and spacers were filtered down to 100 nt after the extension step. For inverted, mirror and direct motifs, the repeat length was filtered to a minimum of 10 nt after extension. Note that 100 nt spacer limits were used to find the largest set of potentially relevant motifs, in order to ensure that flanking regions were devoid of any possible motif, but stricter limits were later used as filters to investigate the role of spacer length.

Potential Z-DNA-forming regions were identified based on two widely-used definitions. The first definition specifies that 10 nt or more must consist of alternating purines and pyrimidines, with the exclusion of any AT dinucleotides. This search was accomplished by replacing all purines and pyrimidines in the reference genome sequence with the symbols ‘R’ and ‘Y’ respectively, and using ‘regex’ to find coordinates matching ‘RYRYRYRY’ motifs. Next, the ‘ATGC’ reference sequence was retrieved for each set of coordinates, in order to filter out motifs containing ‘AT’ dinucleotides. Then, each entry was extended in the 5’ and 3’ directions until each entry no longer contained alternating purines and pyrimidines, contained an AT dinucleotide, or reached a masked region. Finally, entries shorter than 10 nt were filtered out. The second definition specifies G followed by Y for at least 10 nucleotides. The search was performed as above with this additional restriction, as well as for the opposite strand (C followed by R). All entries from the second, more restrictive Z-DNA definition are contained within the first Z-DNA definition.

For G-quadruplex motifs, we began with a set of coordinates derived from experimental mapping of quadruplex-forming structures under two different quadruplex-stabilizing conditions^51^. PDS is a stronger stabilizer, detecting weaker G4 motifs, while K^+^ is a weaker stabilizer, detecting stronger G4 motifs. Data is shown for K^+^ conditions unless otherwise specified. The coordinates were lifted over from hg19 to hg38, reduced to a set of minimal overlapping coordinates, then filtered for those containing at least 4 GGG-tracts (or CCC tracts, depending on the strand). Those coordinates were then trimmed to start and end with GGG-tracts, and filtered to remove any tracts longer than 1kb, as well as any tracts overlapping masked regions.

STR, inverted, mirror, direct, Z-DNA and G4 motif coordinates were combined into a single database, in order to uncover overlaps between motifs. Because STRs were masked prior to other motif searches, STRs with multiple axes of symmetry exist only in the STR category. Inverted, mirror, direct, Z-DNA and G4 motifs fully or partially overlapping any other motif were removed, creating a database of unique repeat categories. We additionally set a buffer zone of 20 nt for repeat motifs and 5 nt for masked regions (Repeatmasker transposons, centromeres, etc.) to avoid any near-overlaps. The buffer zone was initially set to 150 nt (data not shown), related to the typical Illumina read length, but was reduced after we observed no mutational effects extending beyond 20 nt, in order to exclude fewer repeats and gain additional power.

### Flanking mutation frequency calculation and progressive filtering

For each repeat motif, we consider the start and end position of the motif, as well as the distance from the start position to the nearest 5’ motif or masked region, and the distance from the end position to the nearest 3’ motif or masked region, using an additional buffer zone as described above. We thus generate a list of all flanking coordinates that are completely free of the effects of other repetitive regions. The coordinates are used to generate a count of trinucleotides at these positions, as well as a count of any overlapping mutations in the gnomAD rare variant data set. The count of variants is split according to trinucleotide and mutation type (AAA>ACA, AAA>ATA, etc.) Indels were also tabulated according to trinucleotide context, using the trinucleotide of the insertion point and the trinucleotide at the center of the deleted sequence. To combine reverse-complementary STRs, or G-strand with C-strand G4 motifs, 5’ and 3’ coordinates were reversed (ie. position +1 for G strand and position -1 for C strand are combined).

For each position, the trinucleotide mutation counts are divided by the trinucleotide counts multiplied by the number of genomes in the gnomAD database to determine mutation frequency per trinucleotide per genome. This frequency is then normalized to the mutation frequency per trinucleotide per genome at all positions in the genome. By accounting for the expected genome-wide trinucleotide mutation frequency at every position, this eliminates potential artifacts from a non-random composition of nucleotides.

We implemented an additional correction for higher-order G/C content effects on SNVs. We calculated the trinucleotide mutation frequency across the genome in 51 nt overlapping windows, and then adjusted relative mutation frequencies based on this. This window size appears to capture some portion of interactions between extreme GC content regions, such as CpG islands, and mutagenesis/sequencing errors **(Fig S8)**.

The normalized values were then used to create a weighted average mutation frequency per position, with the weights being the trinucleotide mutation counts. To measure mutation frequencies in terms of the relative change from the background mutation rate, we further normalized the weighted average mutation frequency to that derived from a set of ∼100,000 random genomic coordinates (not containing any previously masked regions). Random non-repetitive regions have some baseline mutation frequency under a given QC-filtering regime, and repeat motifs may induce (or protect from) additional mutations beyond this baseline.

For each position, 95% binomial proportion confidence intervals were calculated, with the number of successes being the total number of mutations across all trinucleotides, and the number of trials being the total number of trinucleotides multiplied by the number of genomes in gnomAD. The upper and lower frequencies were then adjusted based on the ratio of the trinucleotide-normalized and weighted mutation frequency to a weighted average of unnormalized trinucleotide frequencies, resulting in normalized weighted confidence intervals.

For mutations surrounding motifs, for clarity we examine perfect motifs, i.e. motifs with no detected interruptions. Additionally, we exclude the bottom 80% of motif lengths for each category. Because shorter motifs are exponentially more common, excluding the bottom 80% by length typically excludes the very shortest motifs in the database, which could dilute a true signal from longer repeats.

### Mutation frequency calculation within motifs, and motif-perfecting mutations

After generating the database of repeat motif coordinates, we searched within each motif for interruptions allowed by the algorithm. For STRs, we separated in-frame and out-of-frame interruptions. In-frame interruptions, which we examine here, can be corrected by mutations, while out-of-frame interruptions can be corrected by indels. For other motifs, we only searched for in-frame interruptions. We measured mutation frequency separately at interrupted sites, and generated separate counts for mutations that restore the perfect motif, and mutations which do not. Mutation frequencies were normalized and weighted as above, with the additional correction that one of three possible mutations can perfect the motif, while two of three cannot.

To estimate the absolute mutation frequency for indels and SNVs within STRs, we multiplied the relative mutation frequencies calculated herein by estimates for absolute SNV and indel frequency per cell per generation, *u*_*bs*_ and *u*_*id*_, as directly measured by Sung, *et al*^52^.

For inverted and mirror repeats, we separately counted mutations that convert the first and last position of the spacer into an extended symmetry, and the same for the first 5’ and 3’ flanking positions. For direct repeats, extending symmetry requires mutation of the position 5’ of the left repeat to match the position 5’ of the right repeat and *vice-versa*, or mutation of the position 3’ of the left repeat to match the position 3’ of the right repeat and *vice-versa* (labeled ‘flank’ and ‘spacer’, respectively, for their position relative to the left repeat).

For G4 motifs, we distinguished positions within G-runs (3 or more Gs in a row) and within spacers. We further distinguished 5’, middle and 3’ positions by their trinucleotide sequence: HGG, GGG, GGH, respectively, for G-runs, and GHH, NNN, HHG, respectively, for spacers, with a further designation for 1 nucleotide GHG spacers (H being A, C or T).

### Calculation of error frequency within duplications

In order to calculate the density of single base errors during synthesis of insertions, it was first necessary to determine the sequence serving as the template for each insertion. For direct repeats, we used regex to find the insertion sequence within the motif and/or spacer sequences, allowing for up to 2 mismatches. For STRs, insertions were first screened for similarity to the STR motif, allowing up to 2 errors (or 1 error when the insertion length is 2 nt), and the in-frame motif of the same length was considered to be the template. The total of all sequences where an unambiguous template for the duplication could be found was then used as the denominator, and the number of mismatches used as the numerator to calculate the error frequency. This was also done in trinucleotide context, in order to compare with the genome-wide SNV rate and determine the mutation spectra.

### Calculation of expansion/contraction bias

Expansions of STRs were identified as described in the above section. All deletions within STRs were considered to be contractions, regardless of whether the deletion was in-frame with the motif. To calculate the bias toward expansions or contractions for each STR motif and at each motif length, we multiplied the frequency of insertions by the total length of all confirmed expansions, then divided this by the frequency of deletions multiplied by the total length of all deletions.

## Data availability

The datasets analysed during the current study are freely available from the gnomAD Consortium (https://gnomad.broadinstitute.org/downloads), the UCSC Genome Browser (https://genome.ucsc.edu), the non-B Database (https://nonb-abcc.ncifcrf.gov/apps/nBMST/default/), and other studies as cited.

## Code availability

The code to perform the analysis in the current study is available in a Github repository (https://github.com/ryanmcggg/nonb_motifs).

## Acknowledgements

We would like to thank Vladimir Seplyarskiy and Evan Koch for their important contributions. The project has been funded by National Institutes of Health grants R35-GM127131, 67 R01-MH101244, U01-HG012009 and R01-HG010372.

## Notes

https://github.com/ryanmcggg/nonb_motifs

### Competing Interest Statement

The authors have declared no competing interest.

### Summary of Updates

Extensive revisions, largely driven by a new analysis of indels.

