## Supplementary Figures for "Mutagenesis at non-B DNA motifs in the human genome: a course correction"

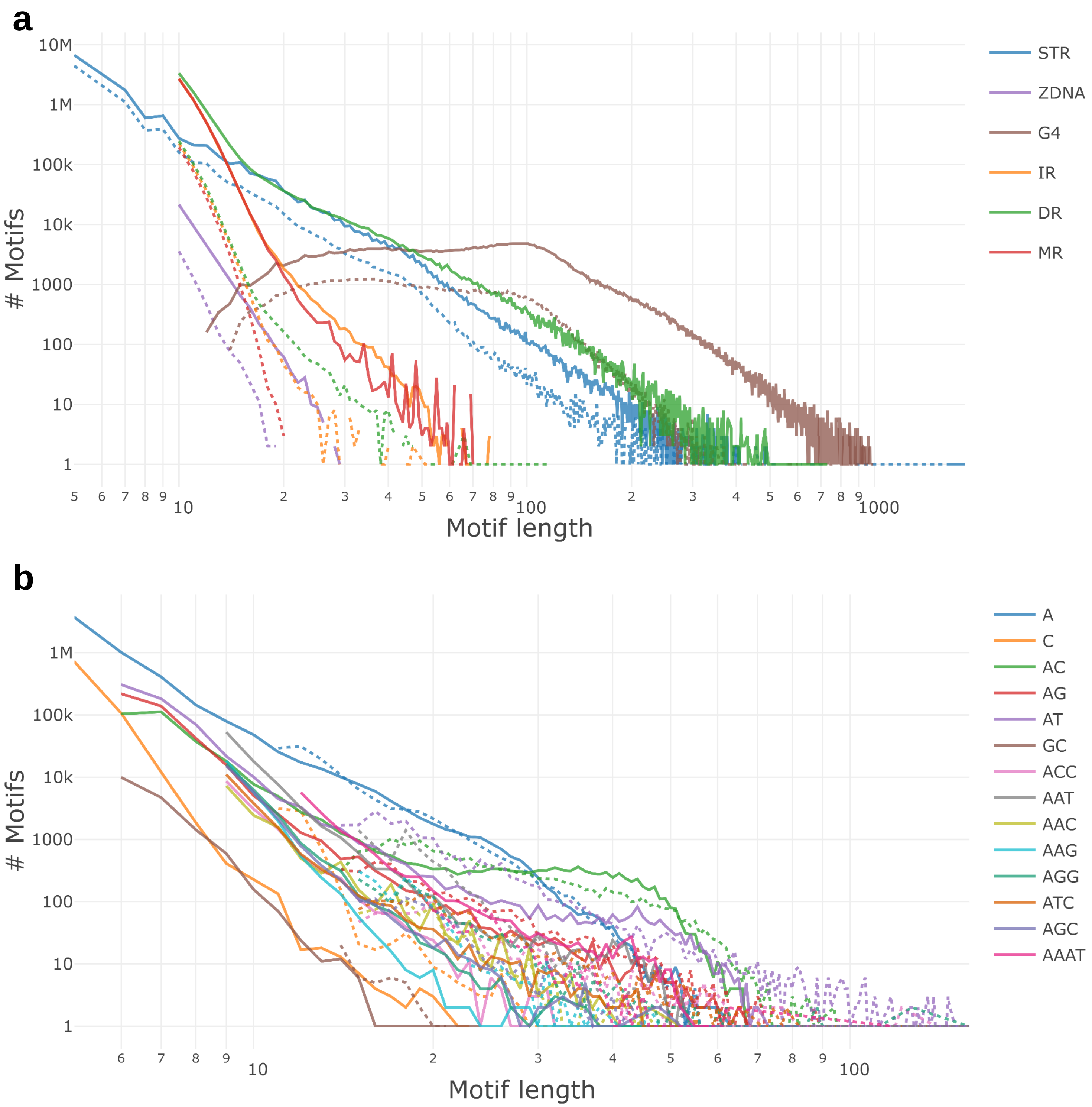


**Fig. S1**: Repeat database overview
**a)** Motif counts per motif length in our custom database. Solid and dashed lines represent counts before and after filtering for uniqueness in the database, respectively. **b)** Motif counts per STR (including reverse-complementary sequences) in our custom database. Solid and dashed lines represent perfect motifs and motifs with in-frame interruptions, respectively.

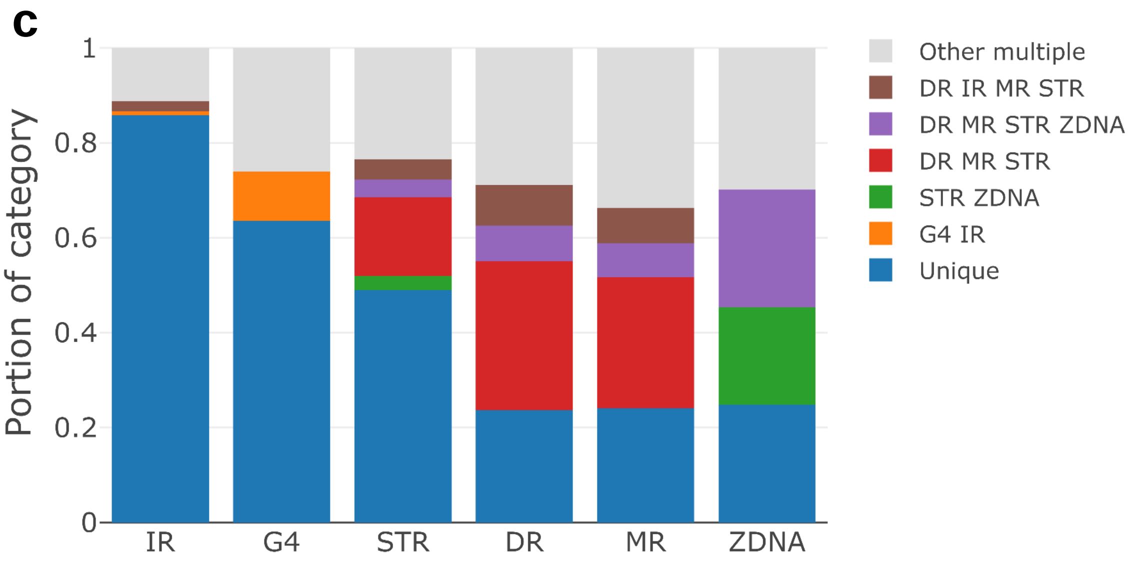


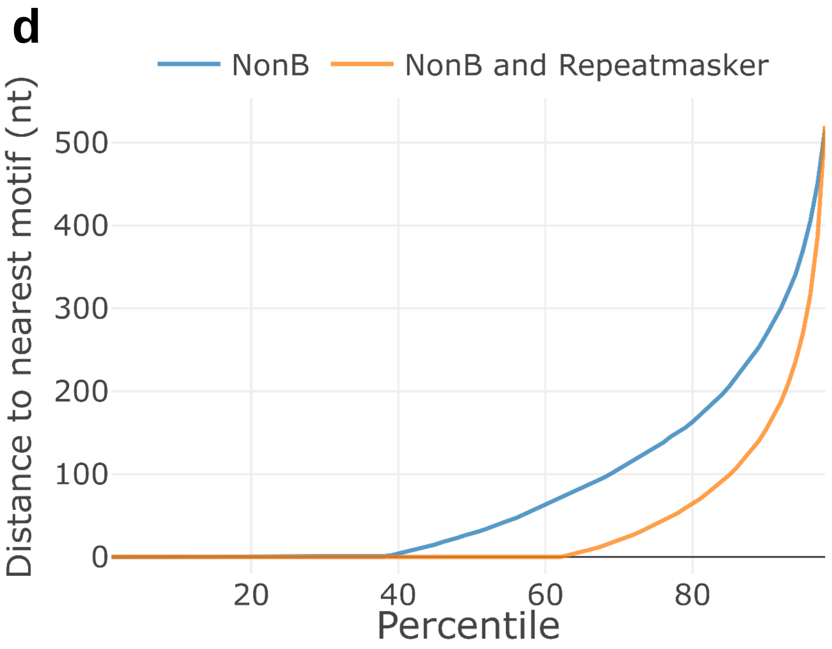


**Fig. S1**: Repeat database overview (continued)
**c)** Overlapping non-B motifs in the "Non-B Database." X-axis indicates categories in Non-B DB. Y-axis indicates the portion of motifs that do not overlap other categories ('unique') and those that overlap additional categories (indicated by color and abbreviation). **d)** Overlapping flanking regions in the "Non-B Database." For each motif in Non-B DB, the distance to the nearest repeat (including transposable elements from Repeatmasker where indicated) on either side was measured, and the smaller of the upstream or downstream values was taken. Overlapping motifs have a distance of 0. X-axis represents the percentile for the range of distances, and the Y-axis indicates the nt distance for each percentile.

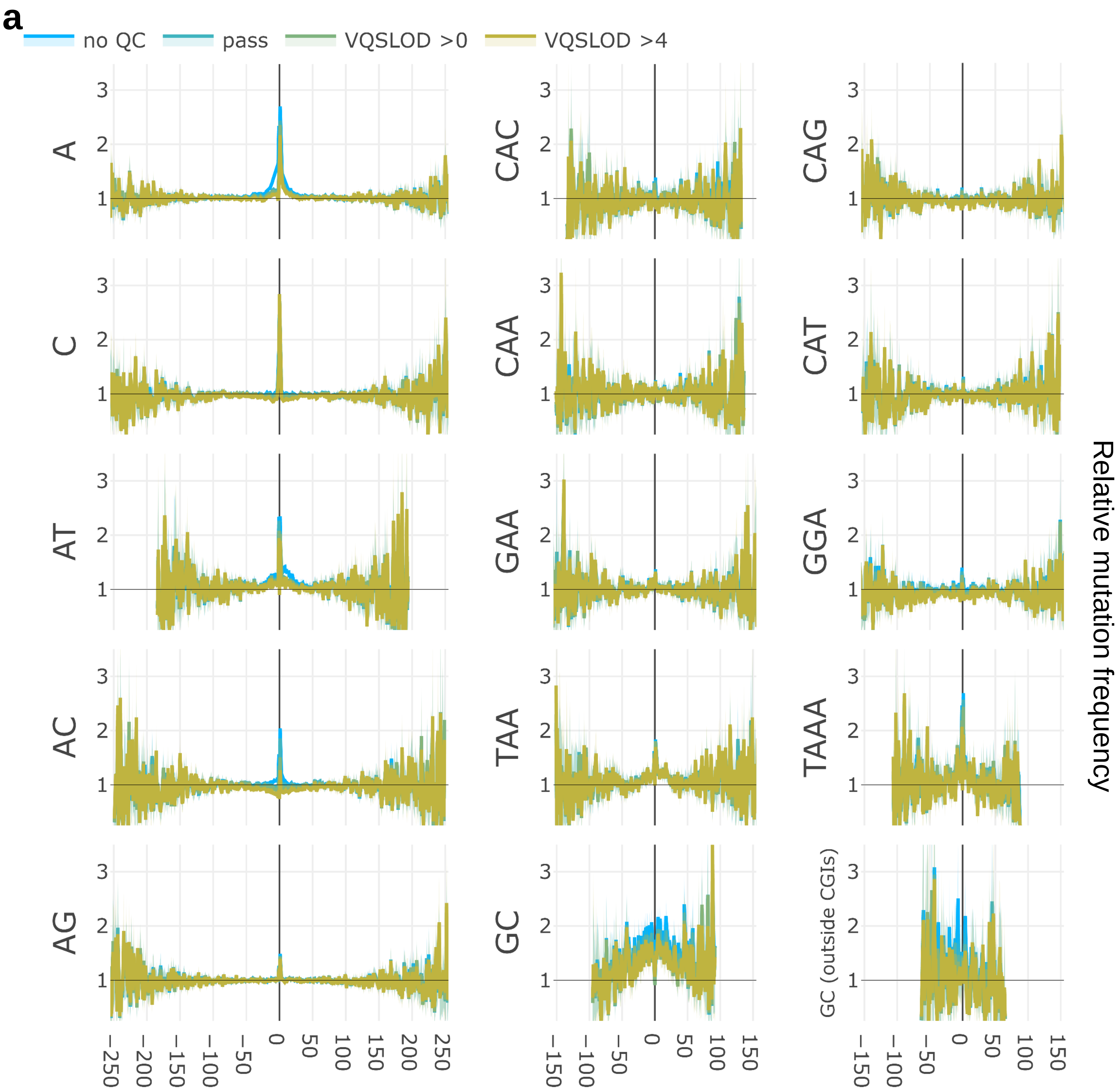


**Fig. S2**: Mutagenesis flanking non-B motifs.
**a)** STR motifs (combined with their reverse complements in a strand-specific manner), filtered to perfect motifs with lengths above the 80% quantile per category. GC dinucleotide repeats shown in the presence or absence of CpG islands (including 2kb shores). X-axes are coordinates relative to central motif (0 represents entirety of the motif). Y-axes are relative mutation frequency. Blue lines: no sequencing quality filters, green and yellow lines: increasingly stringent filters. 95% binomial confidence intervals indicated in transparency.

**b**

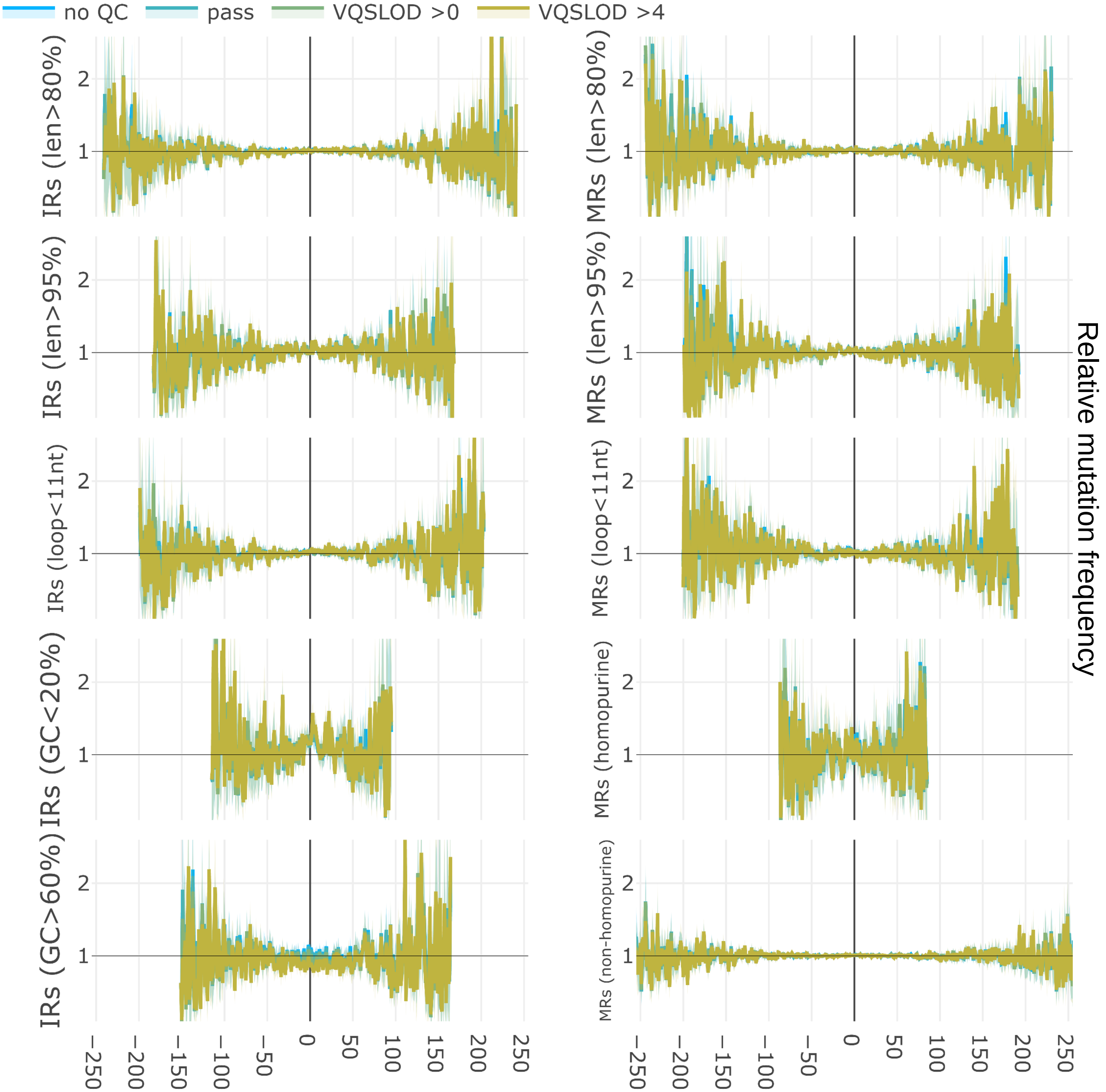


**Fig. S2**: Mutagenesis flanking non-B motifs (continued).
**b)** Inverted (left) and mirror (right) repeats, with additional filters based on length and sequence content, as shown. X-axes are coordinates relative to central motif (0 represents entirety of the motif). Y-axes are relative mutation frequency. Blue lines: no sequencing quality filters, green and yellow lines: increasingly stringent filters. 95% binomial confidence intervals indicated in transparency.

**c**

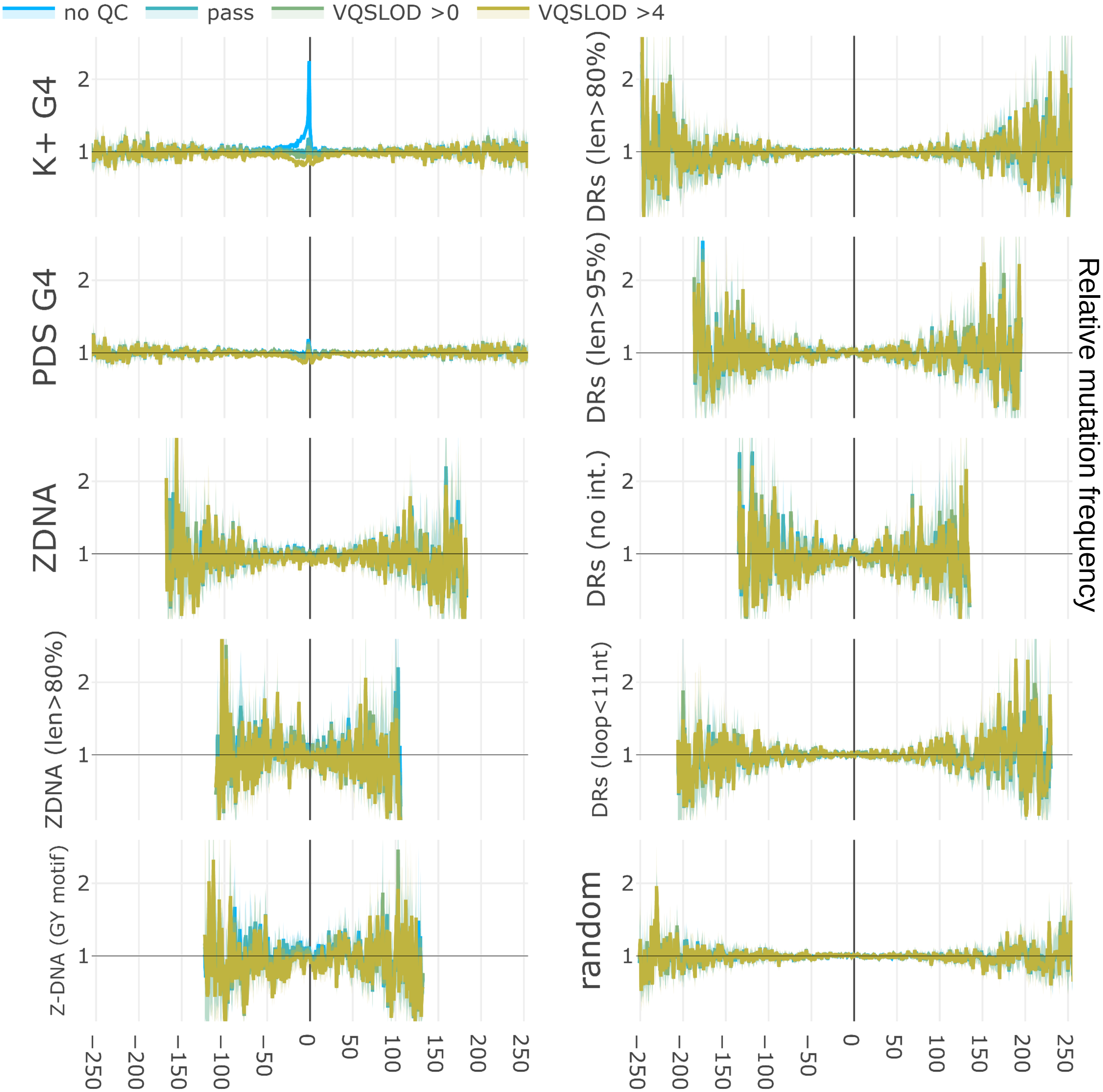


**Fig. S2**: Mutagenesis flanking non-B motifs (continued).
**c)** Additional motifs, including G4 motifs (strand-specific) detected under $K^+$ and PDS conditions, Z-DNA standard and alternate definition requiring GY repeats, direct repeats with length and interruption filters, and random non-repetitive sequences. X-axes are coordinates relative to central motif (0 represents entirety of the motif). Y-axes are relative mutation frequency. Blue lines: no sequencing quality filters, green and yellow lines: increasingly stringent filters. 95% binomial confidence intervals indicated in transparency.

d

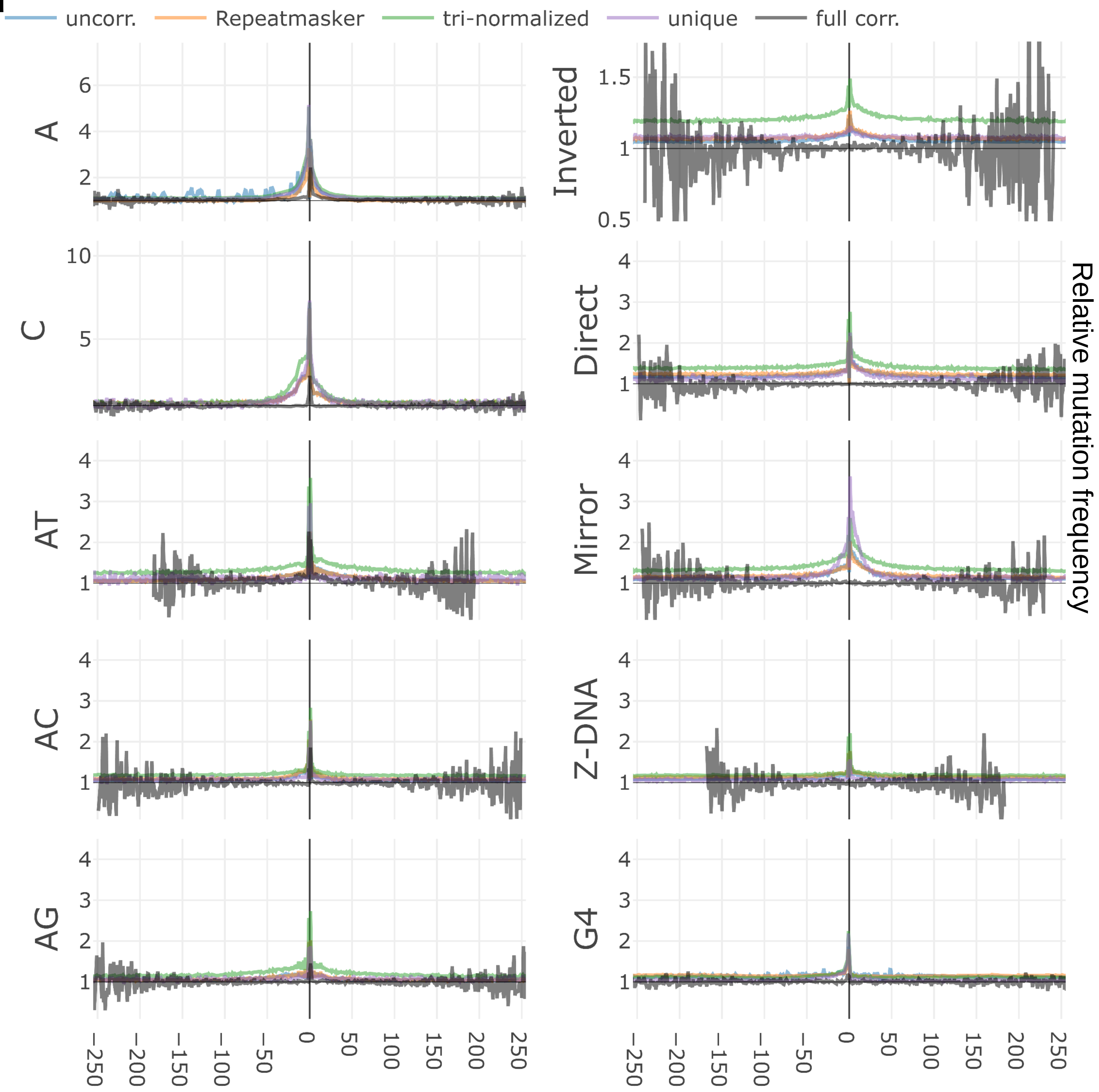


**Fig. S2**: Mutagenesis flanking non-B motifs (continued).
**d)** Analysis with successive layers of corrective measures: blue: no corrections applied to NonB-DB motifs; orange: overlaps with Repeatmasker transposons excluded; green: trinculeotide normalization performed; purple: motif categories filtered for uniqueness; black: flanking regions filtered for presence of any motif, all other corrections applied. X-axes are coordinates relative to central motif (0 represents entirety of the motif). Y-axes are relative mutation frequency.

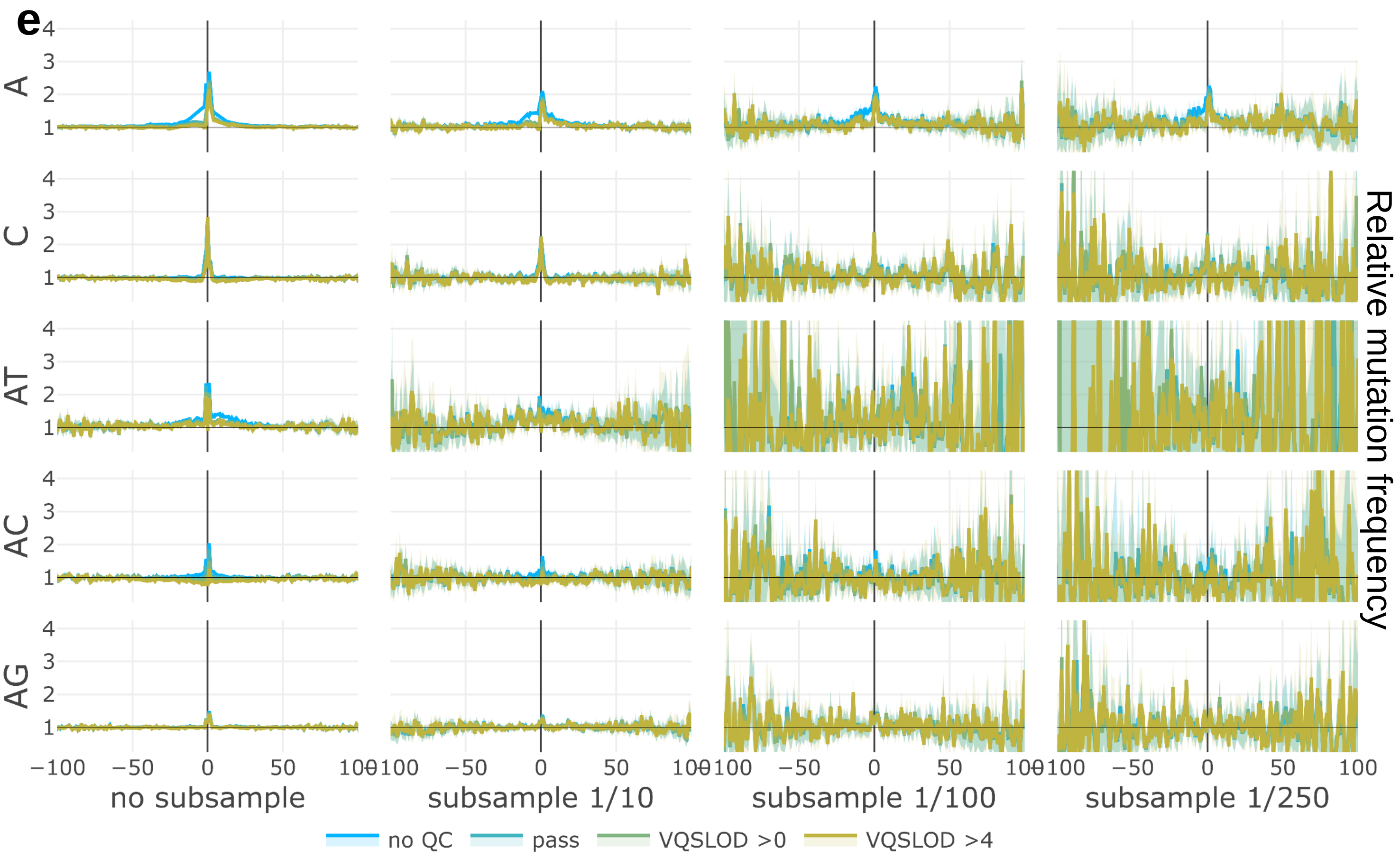


**Fig. S2:** Mutagenesis flanking non-B motifs (continued).
**e)** Relative mutation frequency surrounding STRs, before and after gnomAD subsampling. STR motif (combined with the reverse complementary motif on the opposite strand) indicated on left, subsampling fraction indicated at bottom of each column. X-axes are coordinates relative to motifs, Y-axes are relative mutation frequencies. Blue: no sequencing quality filters, green and yellow: increasingly stringent filters. 95% binomial proportion confidence intervals are presented in transparency.

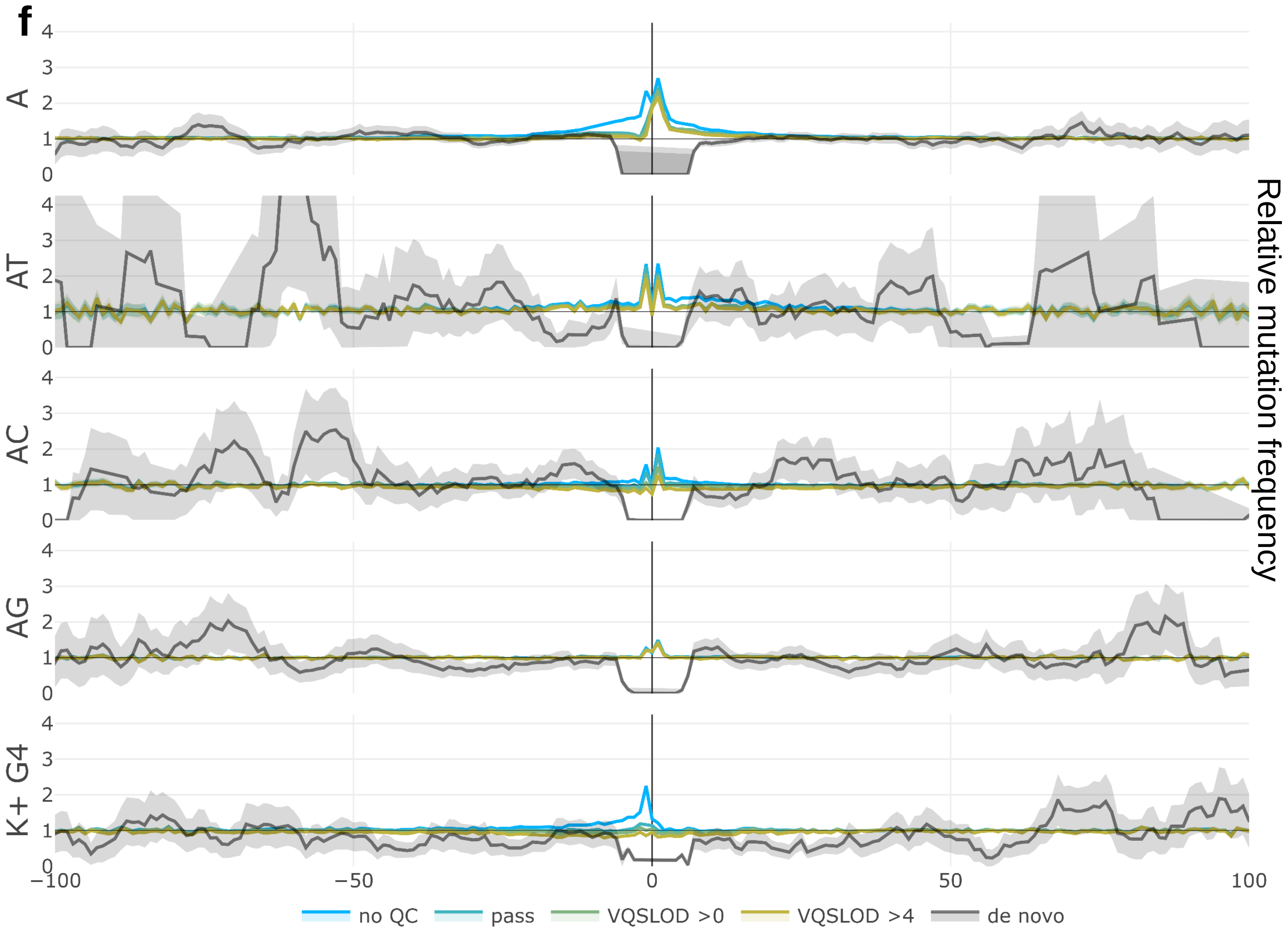


**Fig. S2:** Mutagenesis flanking non-B motifs (continued).
**f)** *De novo* mutagenesis flanking non-B motifs. *De novo* data in gray, gnomAD data shown for comparison. X-axes are coordinates relative to motifs, Y-axes are relative mutation frequencies. 95% binomial proportion confidence intervals are presented in transparency.

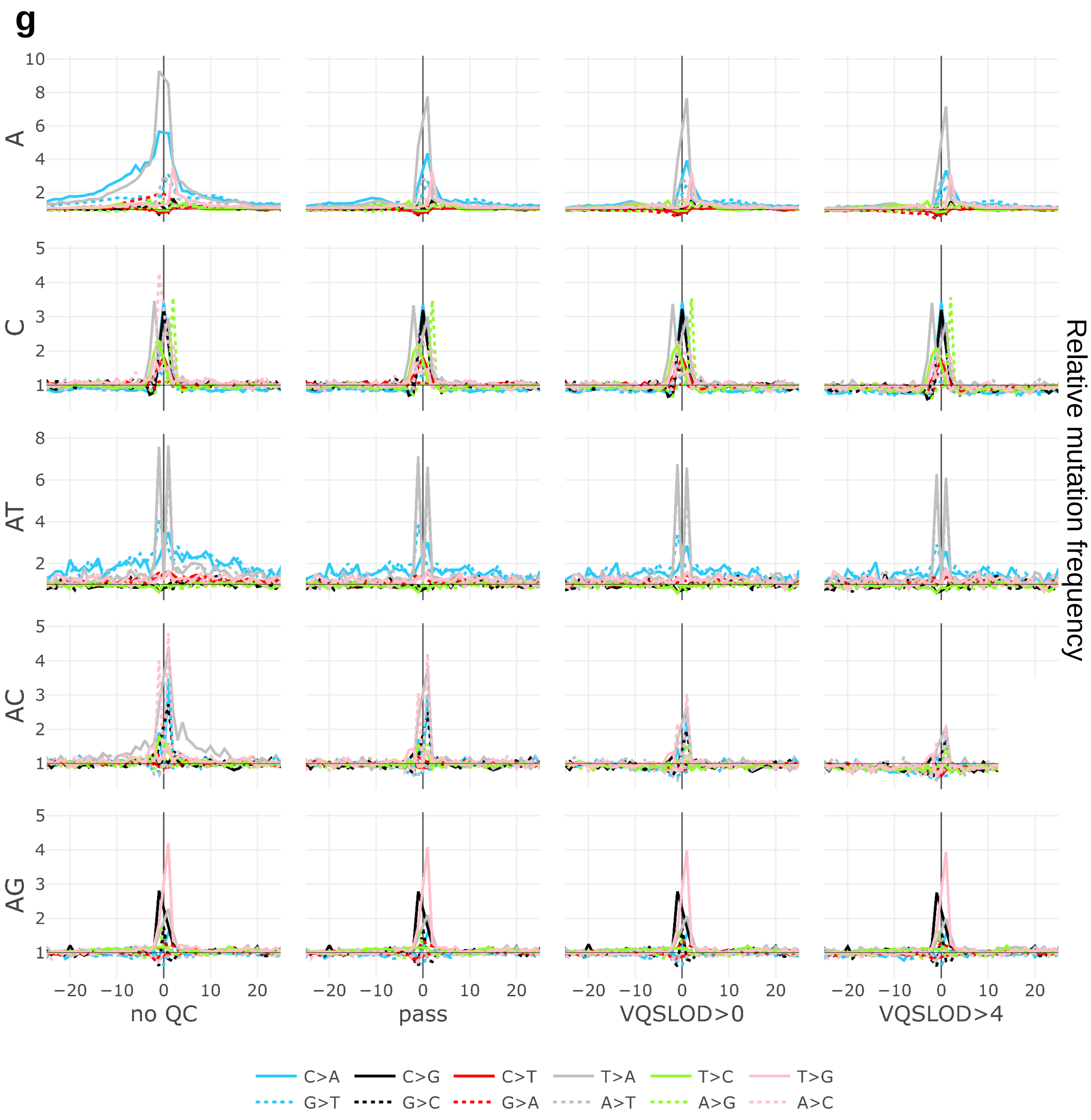


**Fig. S2:** Mutagenesis flanking non-B motifs (continued).

**g)** Relative mutation frequency per-mutation-type surrounding STRs. X-axes are coordinates relative to STRs, Y-axes are relative mutation frequencies. Columns represent the given quality filter. Colors represent different mutation types, and dashed lines represent the reverse-complementary mutation type for each color. For visual clarity, confidence intervals are not presented here. Note that for each motif, the dominant mutation types would generate an extended motif (eg. C>A and T>A mutations would generate extended A-mononucleotide repeats).

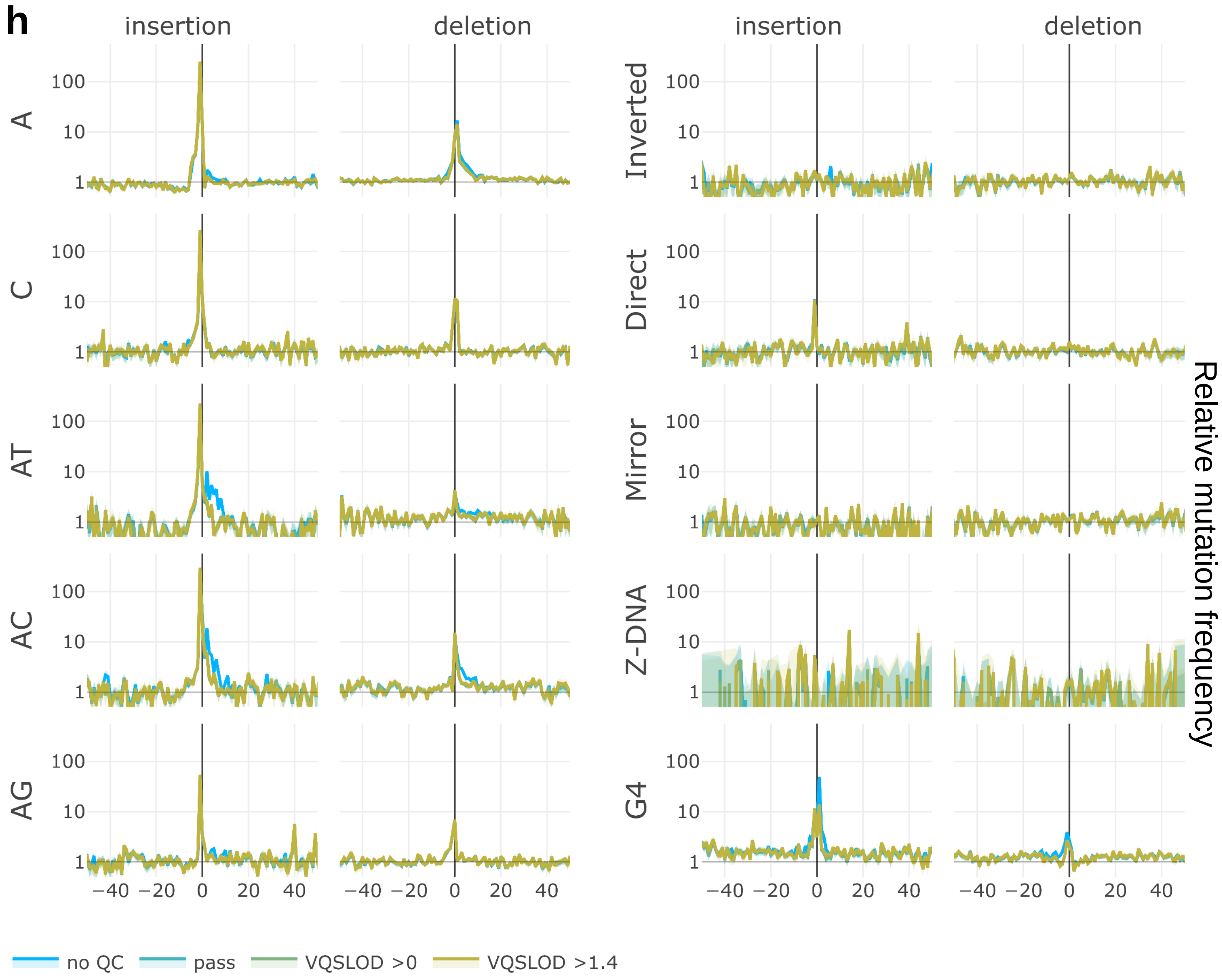


**Fig. S2:** Mutagenesis flanking non-B motifs (continued).
**h)** Insertions and deletions flanking non-B motifs. X-axes are coordinates relative to motifs, Y-axes are relative mutation frequencies. Blue: no sequencing quality filters, green and yellow: increasingly stringent filters. 95% binomial proportion confidence intervals are presented in transparency.

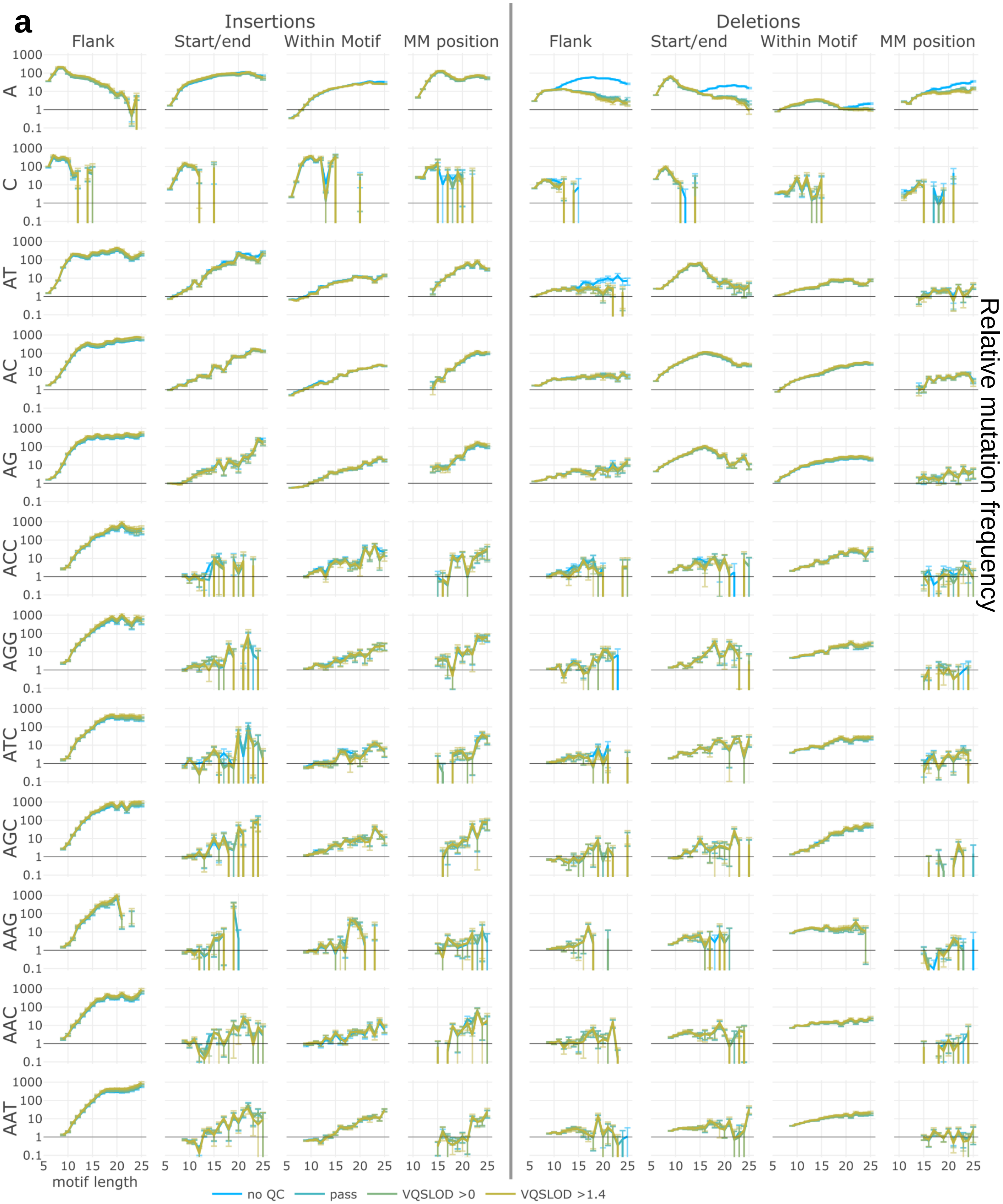


**Fig. S3:** Mutagenesis within non-B motifs.

**a)** Relative indel frequency within STR motifs (and their reverse complements). Blue: no sequencing quality filters, green and yellow: increasingly stringent filters. X-axes are motif lengths. Y-axes are relative mutation frequency. Error bars indicate 95% binomial confidence intervals.

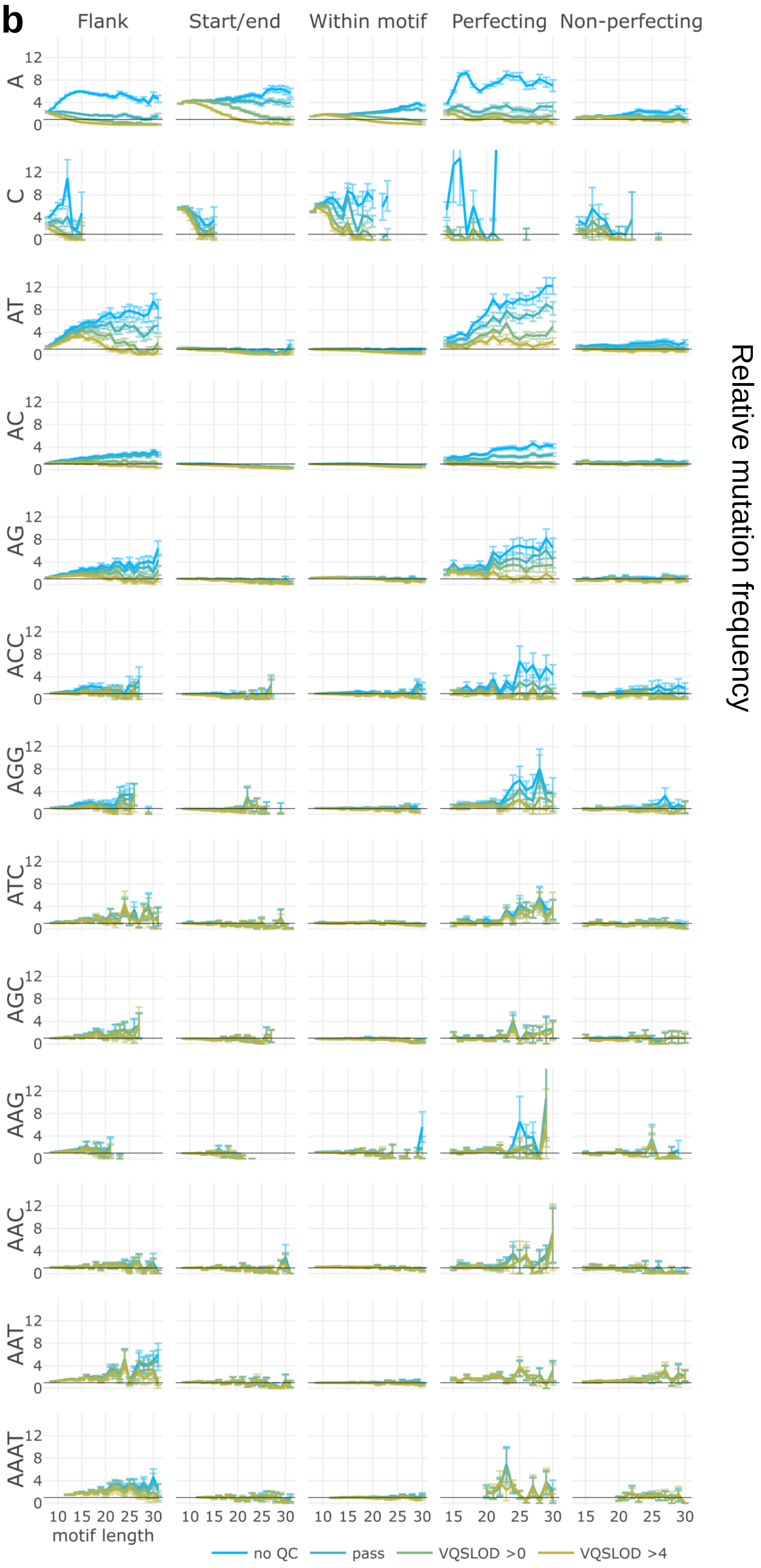


**Fig. S3:** Mutagenesis within non-B motifs (continued). **b)** Relative SNV frequency within STR motifs. Blue: no sequencing quality filters, green and yellow: increasingly stringent filters. X-axes are motif lengths. Y-axes are relative mutation frequency. Error bars indicate 95% binomial confidence intervals.

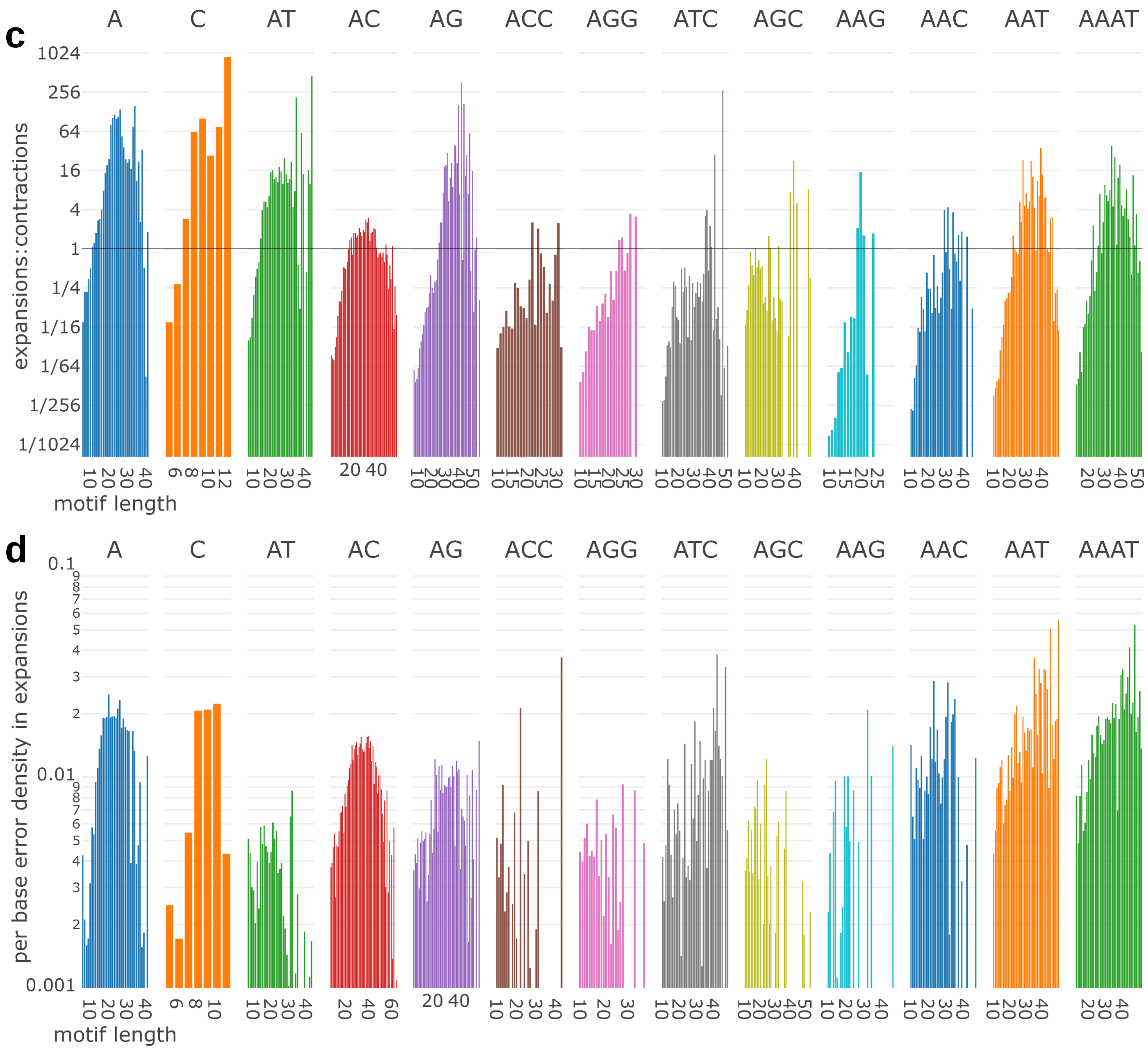


**Fig. S3:** Mutagenesis within non-B motifs (continued).
X axis represents motif length for each STR (combined with its reverse complement). **c)** Ratio of expansions to contractions within STR motifs. **d)** Polymerase error density (single base errors per nucleotide) for expansions within STR motifs.

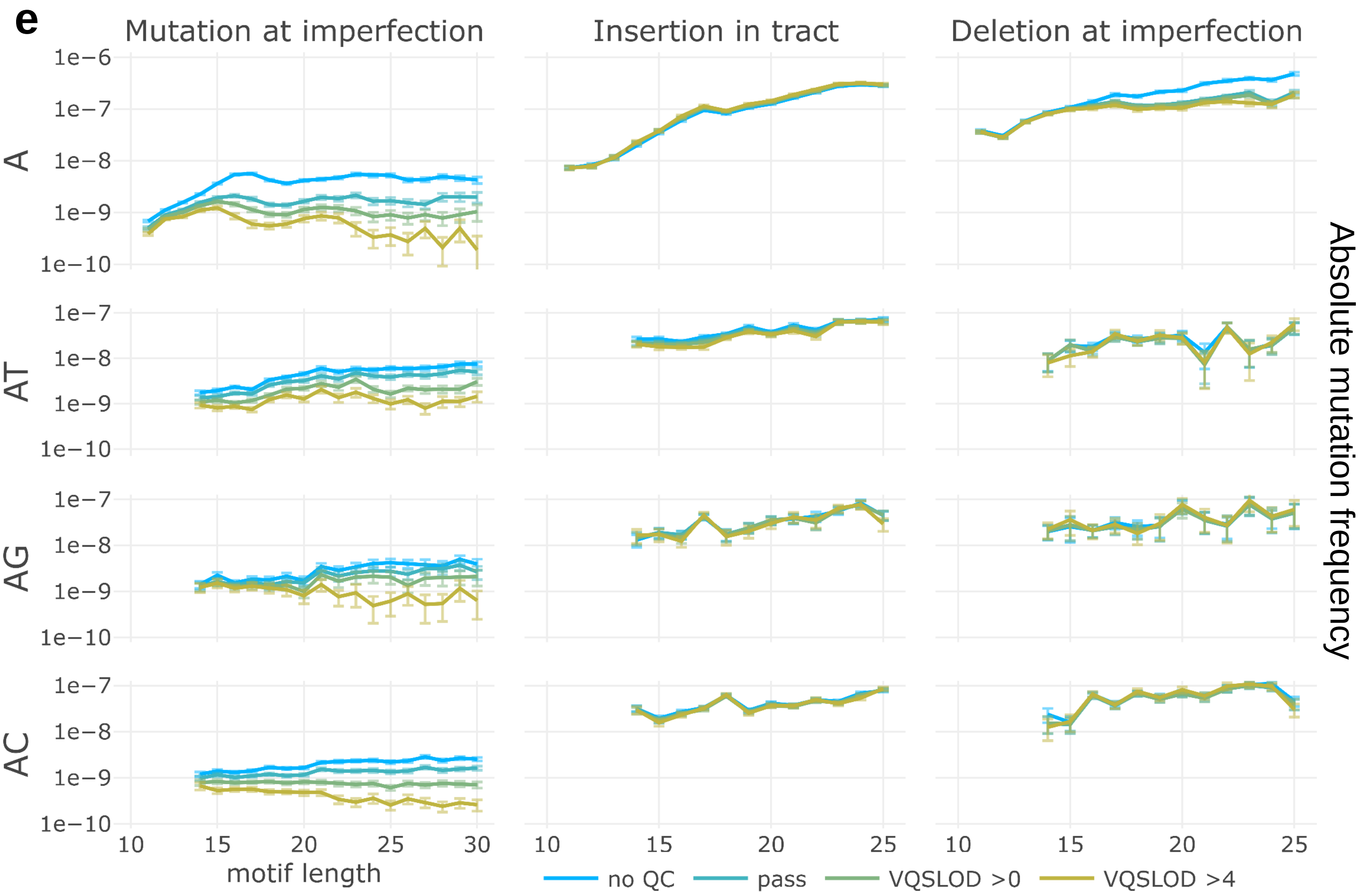


**Fig. S3:** Mutagenesis within non-B motifs (continued).
**e)** Mutation frequencies within STRs, represented as absolute frequencies (mutations per base). Frequency of interruption-perfecting SNVs (left), insertions within imperfect motifs (middle), and deletion of imperfections (right).

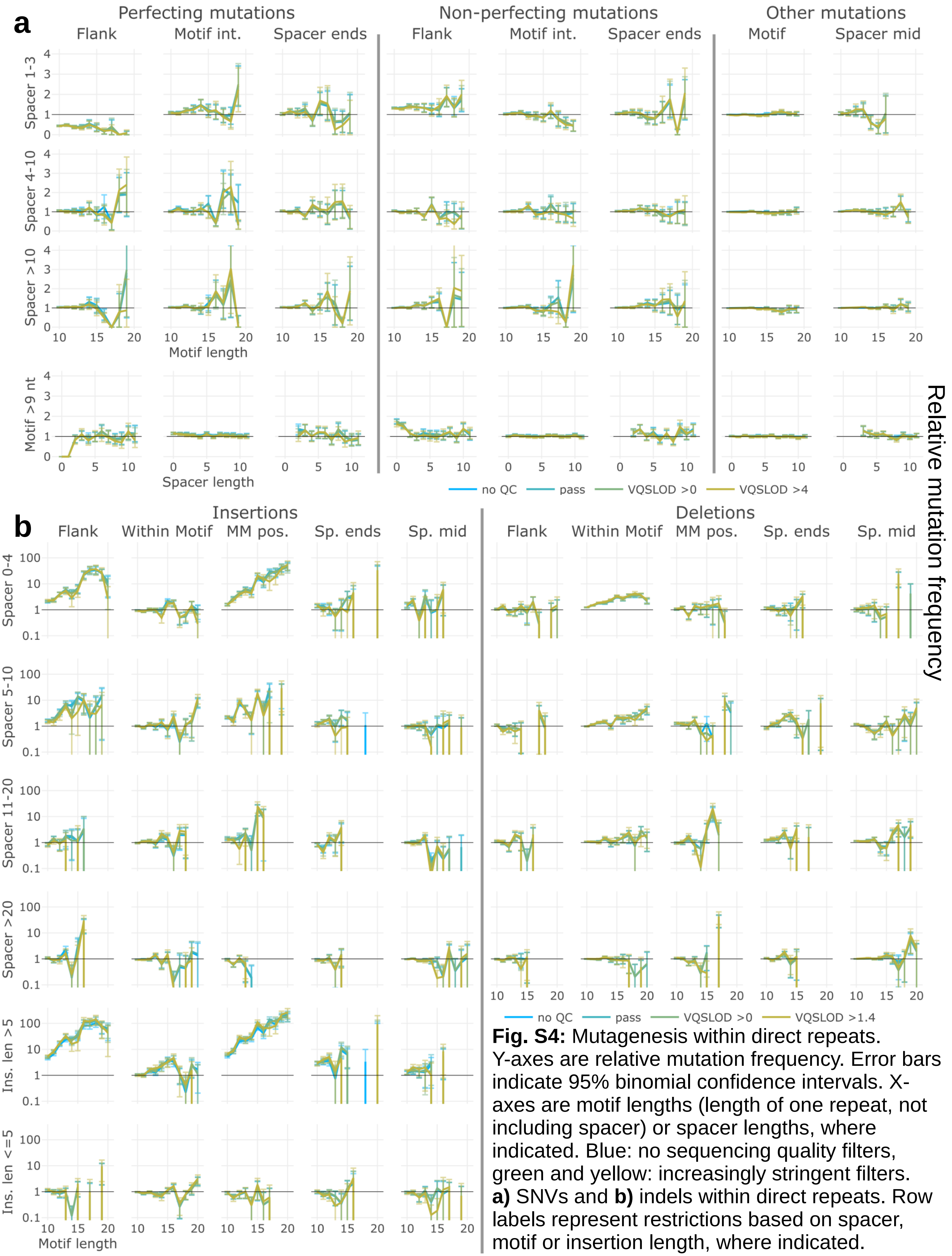


**Fig. S4:** Mutagenesis within direct repeats. Y-axes are relative mutation frequency. Error bars indicate 95% binomial confidence intervals. X-axes are motif lengths (length of one repeat, not including spacer) or spacer lengths, where indicated. Blue: no sequencing quality filters, green and yellow: increasingly stringent filters. **a)** SNVs and **b)** indels within direct repeats. Row labels represent restrictions based on spacer, motif or insertion length, where indicated.

c

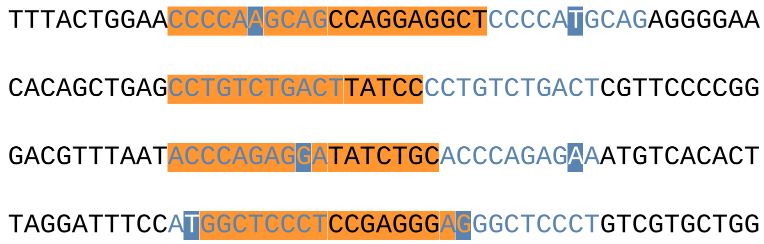


d

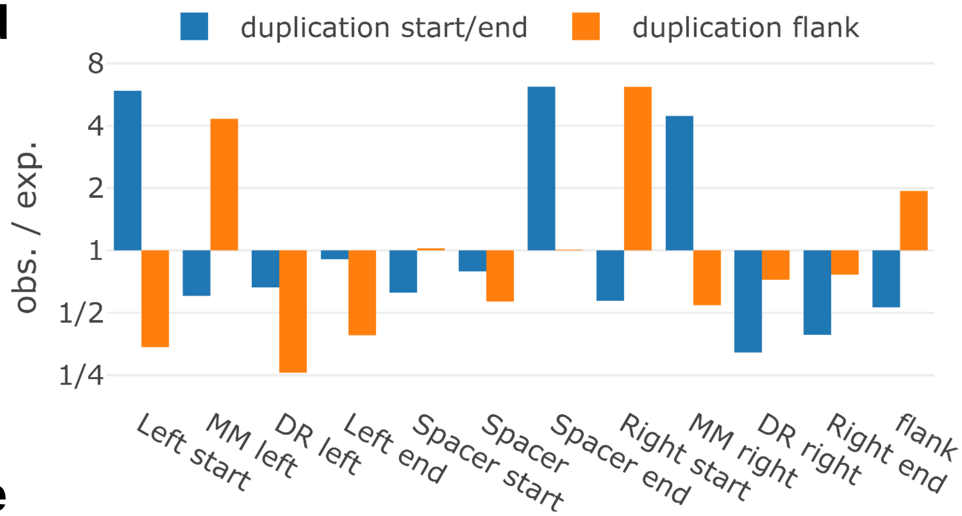


e

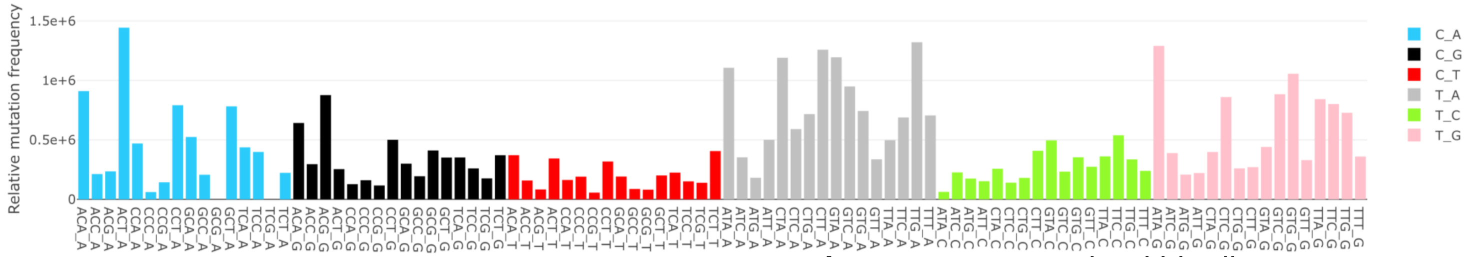


f

| Mut. | Freq. |
|---|---|
| AA>TT | 0.000612 |
| CG>TT | 0.000517 |
| AC>TT | 0.000494 |
| GG>AA | 0.000388 |
| AA>CC | 0.000367 |
| AC>TG | 0.000329 |
| AG>TT | 0.000328 |
| TG>AA | 0.000316 |
| AT>GG | 0.000256 |
| AG>TC | 0.000219 |
| TG>CC | 0.000211 |
| GC>AA | 0.000144 |
| GC>TT | 0.000144 |
| AT>TA | 0.000128 |
| AT>TC | 0.000128 |

g

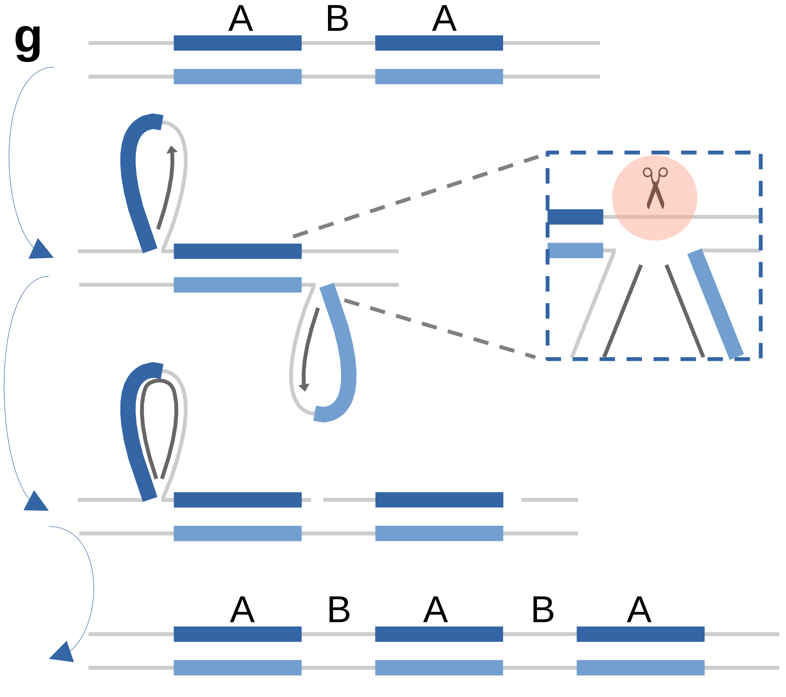


**Fig. S4:** Mutagenesis within direct repeats (continued).
**c)** Examples of duplications within direct repeats. Reference sequence shown. Blue text indicates location of repeat motifs. Blue highlight shows location of motif mismatches. Orange highlight indicates region duplicated in gnomAD. **d)** Duplication position bias. Duplication start/end positions (blue) or positions flanking duplications (orange), categorized their location (ie. start of left motif, mismatch position in left motif, end of spacer, etc.). Frequency of the duplication at each position (observed) is divided by the portion of the motif for which the position accounts (expected). **e)** Spectrum of errors within duplications. X axis represents 96 trinucleotide mutation types. Y axis is the relative increase compared to the frequency of *de novo* mutations per mutation type. **f)** Observed double-nucleotide error frequency within duplications (based on the total number of dinucleotides across all duplication sequences). **g)** Gap repair model explaining direct repeat duplications. The initial A-B-A pattern forms a slipped-strand structure. Post-replicative gap-filling (Polζ) fills in single-stranded loop regions, resulting in 4-way pseudo-Holliday junctions. Cleavage of the top strand allows the filled-in sequence to be ligated into the top strand. Either this can be repeated for the other loop, or replication of the top strand will produce a daughter cell with the A-B-A-B-A pattern.

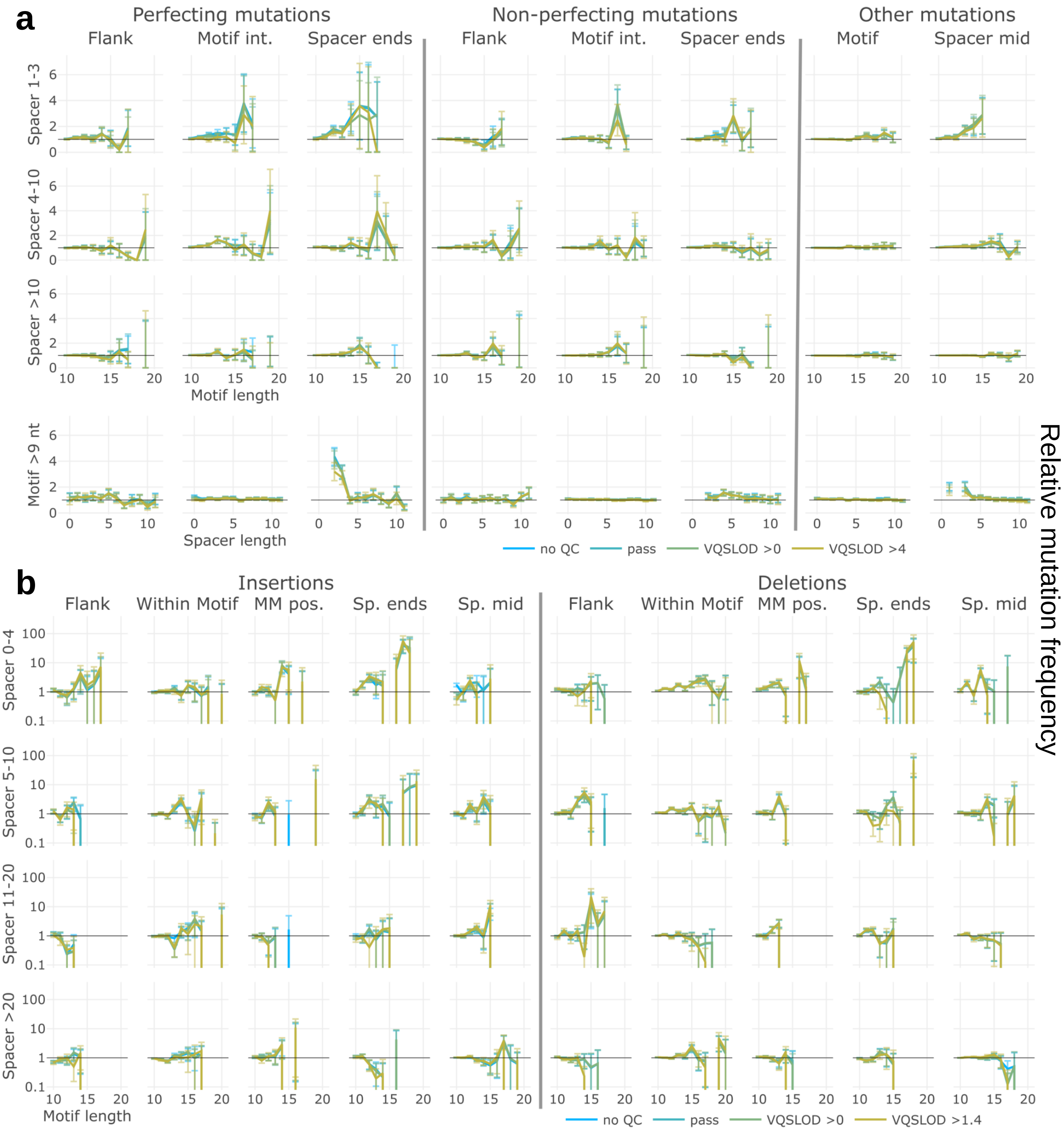


**Fig. S5:** Mutagenesis within inverted repeats.
Y-axes are relative mutation frequency. Error bars indicate 95% binomial confidence intervals. X-axes are motif lengths (length of one repeat, not including spacer) or spacer lengths, where indicated. Blue: no sequencing quality filters, green and yellow: increasingly stringent filters.
**a)** SNVs and **b)** indels within inverted repeats. Row labels represent restrictions based on spacer, motif or insertion length, where indicated.

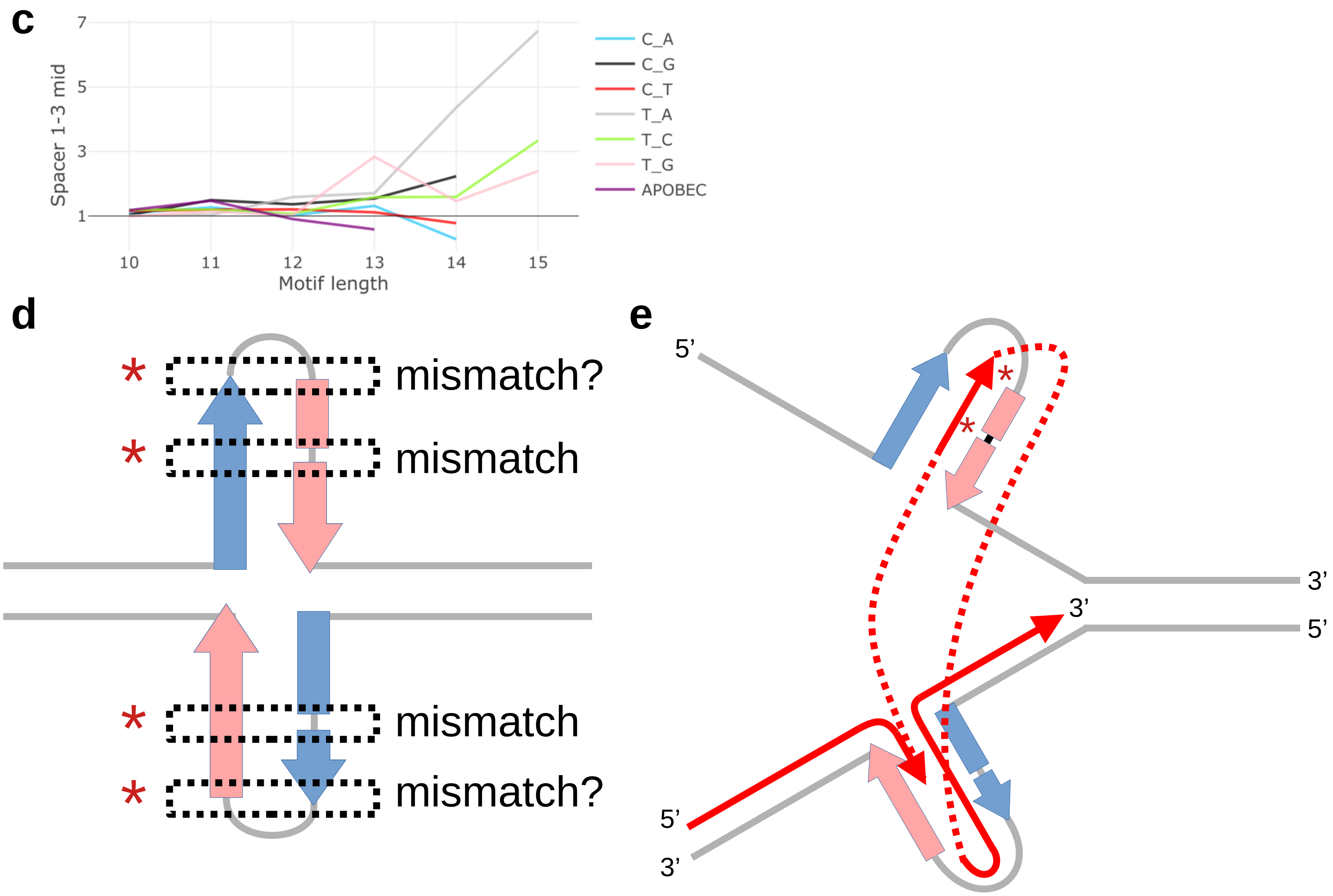


**Fig. S5:** Mutagenesis within inverted repeats (continued).
**c)** Rate of mutations per-mutation type for mutations in the middle of spacer sequences, restricted to 1 and 3 nt spacers (iteration on data in the top right panel of Fig. S4A). Y-axis is relative mutation frequency. X-axis is motif length (length of one repeat, not including spacer). Colors indicate mutation type. “APOBEC” mutations consist of TCN>T mutations. **d)** Mismatch repair model for interruption-perfecting SNVs. * indicates positions of frequent perfecting/extending mutations. Inner imperfections may appear as typical mismatches when the motif forms a stable hairpin/cruciform structure. Loops with 2-3 nt length might also be recognized as mismatches. **e)** Template-switch during replication produces motif-perfecting and inward-extending mutations. The red line follows the path of the polymerase as it proceeds 5’ to 3’. Dotted lines indicate template-switches. * indicates positions of frequent perfecting/extending mutations.

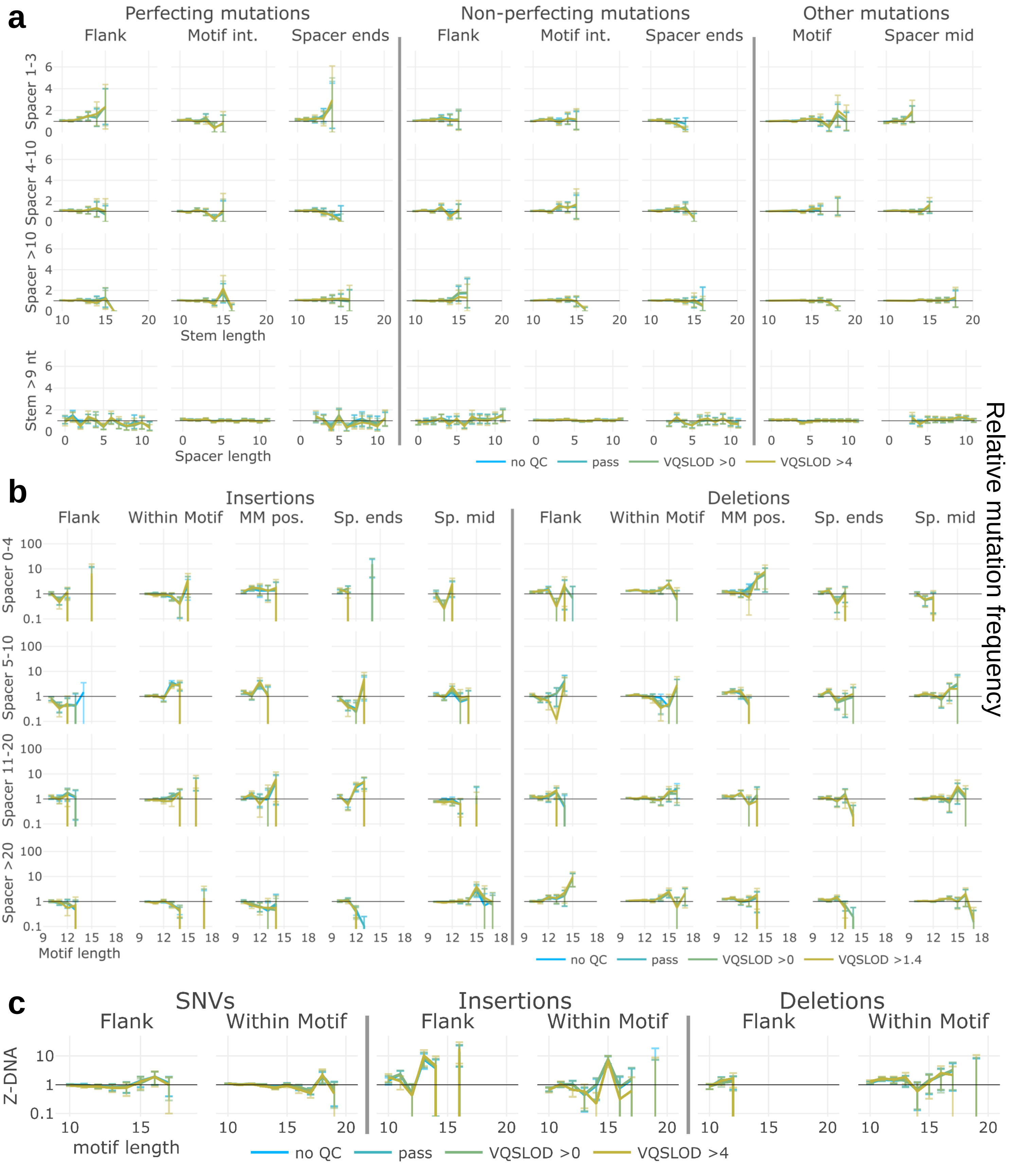


**Fig. S6**: Mutations within mirror repeats and Z-DNA motifs
Y-axes are relative mutation frequency. Error bars indicate 95% binomial confidence intervals. X-axes are motif lengths (length of one repeat, not including spacer) or spacer lengths, where indicated. Blue: no sequencing quality filters, green and yellow: increasingly stringent filters.
**a)** SNVs and **b)** indels within mirror repeats. Row labels represent restrictions based on spacer, motif or insertion length, where indicated. **c)** SNVs and indels within Z-DNA motifs

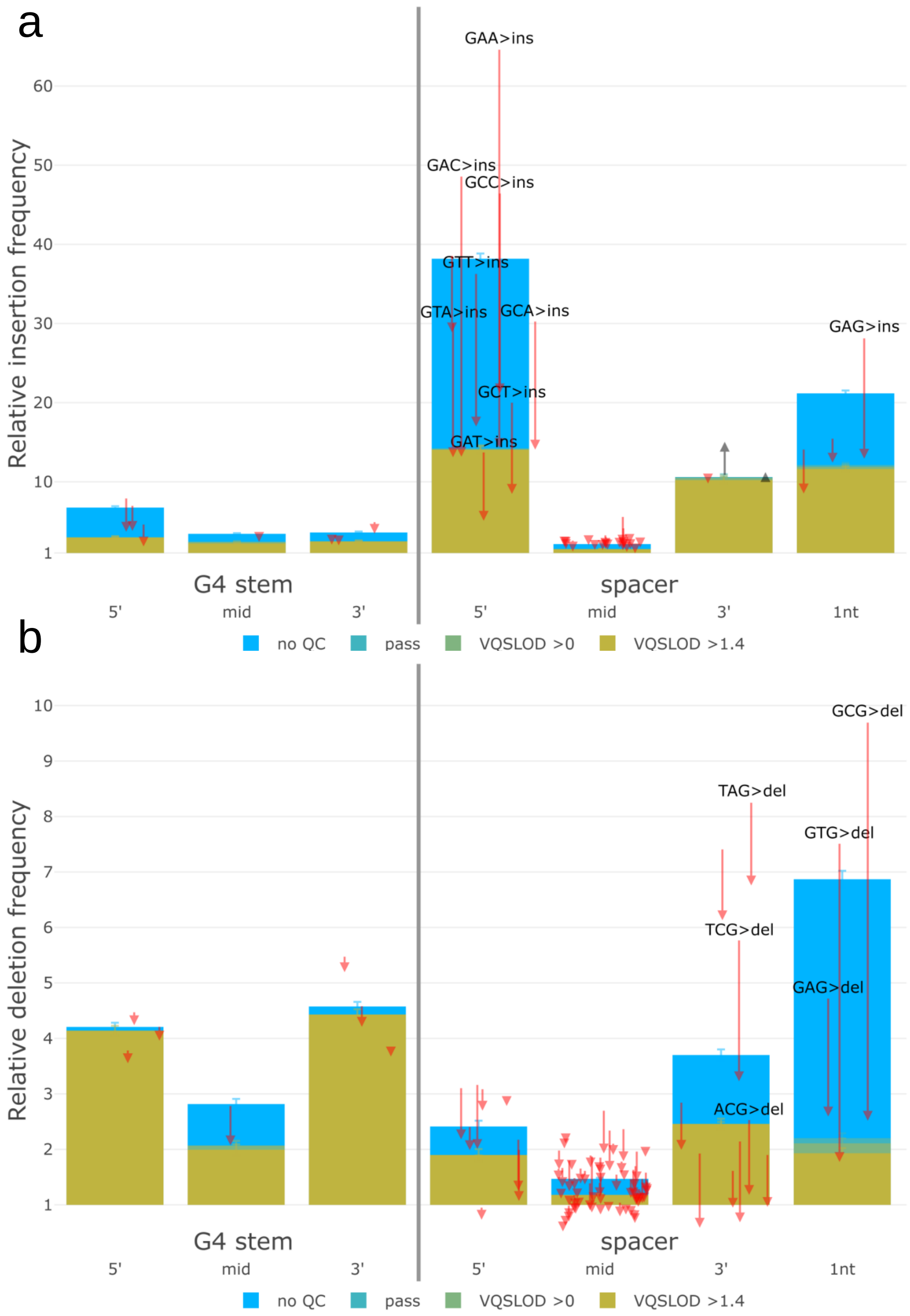


**Fig. S7**: G4 motifs are prone to recurrent sequencing errors. **a)** Insertions and **b)** deletions within G4 motifs. X-axis indicates positions within G-runs, or spacers between G-runs. Spacers of 1 nt in length are categorized separately. Y-axis is relative mutation frequency. Blue: no sequencing quality filters, green and yellow: increasingly stringent filters. Error bars indicate 95% binomial confidence intervals. Arrows indicate magnitude and direction of change for individual trinucleotide mutation frequencies before and after sequencing quality filters. Mutations with large magnitude changes are highlighted in text.

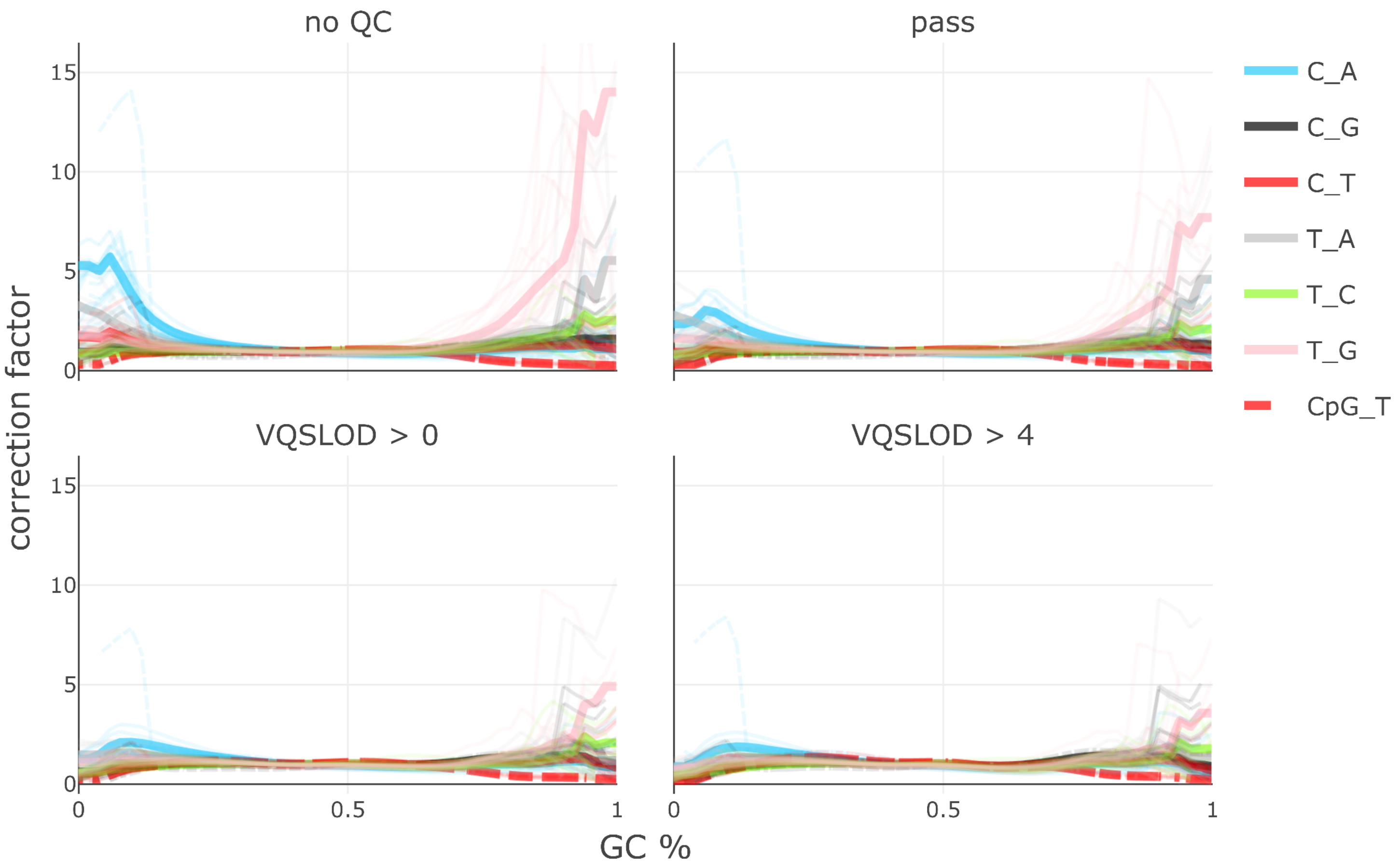


**Fig. S8**: SNV mutation frequency correction based on 51 nt window GC content
X-axis is GC content measured +/- 25 nt from each SNV. Y-axis is the correction factor applied to mutation frequency. Colors represent mutation types. Thin bars represent individual trinucleotide mutation types (eg. ACA>ATA), and thick bars represent median per mutation type. Dashed lines represent NCG trinucleotide contexts, eg. CpG contexts. Note that the expected CpG transition rate in GC-rich regions is elevated due to deamination of methyl-CpGs, and therefore the correction factor is < 1.
